# The Arp2/3 complex promotes periodic removal of Pak1-mediated negative feedback to facilitate anticorrelated Cdc42 oscillations

**DOI:** 10.1101/2023.11.08.566261

**Authors:** Marcus Harrell, Ziyi Liu, Bethany F Campbell, Olivia Chinsen, Tian Hong, Maitreyi Das

## Abstract

The conserved GTPase Cdc42 is a major regulator of polarized growth in most eukaryotes. Cdc42 periodically cycles between active and inactive states at sites of polarized growth. These periodic cycles are caused by positive feedback and time-delayed negative feedback loops. In the bipolar yeast *S. pombe*, both growing ends must regulate Cdc42 activity. At each cell end, Cdc42 activity recruits the Pak1 kinase which prevents further Cdc42 activation thus establishing negative feedback. It is unclear how Cdc42 activation returns to the end after Pak1-dependent negative feedback. Using genetic and chemical perturbations, we find that disrupting branched actin-mediated endocytosis disables Cdc42 reactivation at the cell ends. With our experimental data and mathematical models, we show that endocytosis-dependent Pak1 removal from the cell ends allows the Cdc42 activator Scd1 to return to that end to enable reactivation of Cdc42. Moreover, we show that Pak1 elicits its own removal via activation of endocytosis. In agreement with these observations, our model and experimental data show that in each oscillatory cycle, Cdc42 activation increases followed by an increase in Pak1 recruitment at that end. These findings provide a deeper insight into the self-organization of Cdc42 regulation and reveal previously unknown feedback with endocytosis in the establishment of cell polarity.

## INTRODUCTION

The relationship between structure and function is a core tenet of biology. This holds true for cells, which must achieve a specific shape for their function. As such, the plethora of environments and purposes that cells occupy results in a wide diversity of cellular shapes and corresponding functions. These diverse shapes are attributed to polarized growth, wherein cells organize the cytoskeleton to promote growth from specific sites [1, 2]. While several works have investigated how cells polarize and acquire shape, the fundamentals of polarization are still not well understood. This is mainly because polarization involves multiple signaling proteins and pathways that are tightly regulated. Major regulators of polarized growth across most eukaryotes are the highly conserved Rho Family GTPases which consists of Cdc42, Rac, and Rho[3]. These GTPases are tightly regulated to promote cell motility, cell shape, and proliferation [3–5]. For example, the GTPases spatiotemporally regulate cell protrusions, speed and direction during migration, and promote different aspects of cell differentiation in synapse formation [6–8]. These findings indicate that the GTPases are regulated via complex higher-order pathways to ensure their proper spatiotemporal activation. Given the complexity of these pathways and the limitations of the mammalian systems, the molecular mechanisms governing cell polarization are not well understood.

The fission yeast, *Schizosaccharomyces pombe* grows in a bipolar manner from the two cell ends. In fission yeast growth initiates at the old end, the end that existed in the previous generation, upon completion of cell division. As the cell reaches a certain size in the G2 phase, the new end that is formed as a result of cell division also initiates growth resulting in a bipolar growth pattern [9]. Similar to most eukaryotes, Cdc42 is a major regulator of cell polarization in fission yeast [10]. Fission yeast share the same conserved mechanisms as higher eukaryotes making it an excellent model to understand higher-order molecular pathways that promote polarization and bipolarity. During bipolar growth in fission yeast, Cdc42 activity cycles on and off in a periodic manner often resulting in oscillations [11]. Cdc42 oscillations are dictated by cycles of positive feedback and time-delayed negative feedback via its regulators resulting in bipolar growth [11–13]. Cdc42 activation occur in an anti-correlated manner between the two growing ends suggesting that the growing ends must compete for resources to sustain Cdc42 activity and growth [11, 13–16]. Due to this competition, Cdc42 must be inactivated at one cell-end for the opposite end to activate Cdc42 and vice-versa [11, 16]. Disruption of these Cdc42 activation cycles leads to a loss of bipolar growth [11]. It is not known how Cdc42 activity periodically returns to a cell end after each cycle of the negative-feedback to allow for the oscillations.

Cdc42 positive-feedback is mediated by the guanine nucleotide exchange factor (GEF) Scd1 and its scaffold protein Scd2 [13, 17, 18]. The scaffold Scd2 binds active Cdc42 and is thus recruited to the growing cell ends [19–21]. Scd2 at the cell ends then recruits Scd1 for further Cdc42 activation thus establishing a positive-feedback. Active Cdc42 can be inactivated by its GTPase activating proteins (GAPs) Rga4, Rga6, and Rga3 [22–25]. As Cdc42 activity reaches a certain threshold it triggers time-delayed negative feedback mediated by the Pak1 kinase [11, 26, 27]. The Pak1 kinase (p21-activated kinase) is a serine/threonine kinase and is highly conserved in regulating growth in most eukaryotes [5, 28]. The Pak1 kinase is activated and recruited to the growing cell ends when it binds active Cdc42 [29, 30]. When active, Pak1 phosphorylates effector proteins to modulate a host of biological pathways [11, 29–34]. In the absence of Pak1 activity, Scd1 and Scd2 accumulate at the cell ends leading to increased Cdc42 activity [11]. In budding yeast, the Pak1 homolog Cla4 has been shown to phosphorylate Cdc24, the Scd1 homolog, to prevent its interaction with Bem1, the Scd2 homolog [35, 36].

Cdc42 regulates multiple pathways such as membrane trafficking and cytoskeletal F-actin organization [3, 4, 10, 37–42]. When activated Cdc42 in turn activates the formin, For3, to nucleate linear actin cables [40, 43–45]. Active Cdc42 also triggers branched actin polymerization, necessary for endocytosis in yeasts, through the WASP protein, Wsp1, and the type 1 myosin, Myo1 [38–40, 44, 46–49] . Both Wsp1 and Myo1 regulate endocytosis by binding the Arp2/3 complex to promote the nucleation of branched actin [47]. We find that Cdc42 regulates endocytosis in fission yeast both at the division site and also at the site of cell growth [42, 48, 50, 51]. However, it is not well understood how membrane trafficking events in turn regulate Cdc42 dynamics.

Using *in vivo* experiments alongside mathematical modeling, we find that branched actin is required for proper removal of Pak1, to allow anticorrelated oscillations. When the branched actin nucleator Arp2/3 complex is inhibited, Pak1 stabilizes at a cell end, preventing localization of the GEF, Scd1, and disrupting the positive and negative feedback loops needed for periodic Cdc42 activity at that end. Arp2/3 complex mediated branched actin is required for endocytosis in fission yeast. While Cdc42 has been shown to regulate endocytosis and actin organization, our findings provide evidence that endocytosis also directly impacts Cdc42 activity [37, 52]. It is widely acknowledged that proper protein localization is vital for activity. Here we show that timely removal of inhibition plays an equally critical role and sets the stage for positive regulators to return and continue the cycles of activity.

## RESULTS

### The Arp2/3 complex is required for anticorrelated oscillations between growing cell ends

There are two F-actin structures in *S. pombe* during interphase: linear actin cables and branched actin networks [53]. Actin cables transverse the length of the cell, physically connecting both ends, and are required for actin mediated delivery and exocytosis [53]. Branched actin patches are primarily located at the cell ends and are required for endocytosis [40, 48]. Given the role for F-actin in membrane trafficking and polarization, we asked if actin structures also regulated Cdc42 anticorrelated oscillations. To disrupt F-actin we treated cells with the inhibitor of actin polymerization Latrunculin A (Lat-A, Fig.1A). We used the fluorescent probe CRIB-3xGFP to specifically visualize active Cdc42 for our experiments [25]. We quantified the competition between the ends within a cell by measuring the correlation of Cdc42 activity between the ends under all experimental conditions (Fig.1B). As has been reported before, we find that Cdc42 oscillations were disrupted in Lat-A treated cells, resulting in loss of CRIB-3xGFP signal at the cell ends and increased depolarized signal along the cell sides [54, 55]. Lat-A treated cells displayed enhanced correlation of CRIB-3xGFP signal between cells ends (Fig.1B).

**Figure 1.**
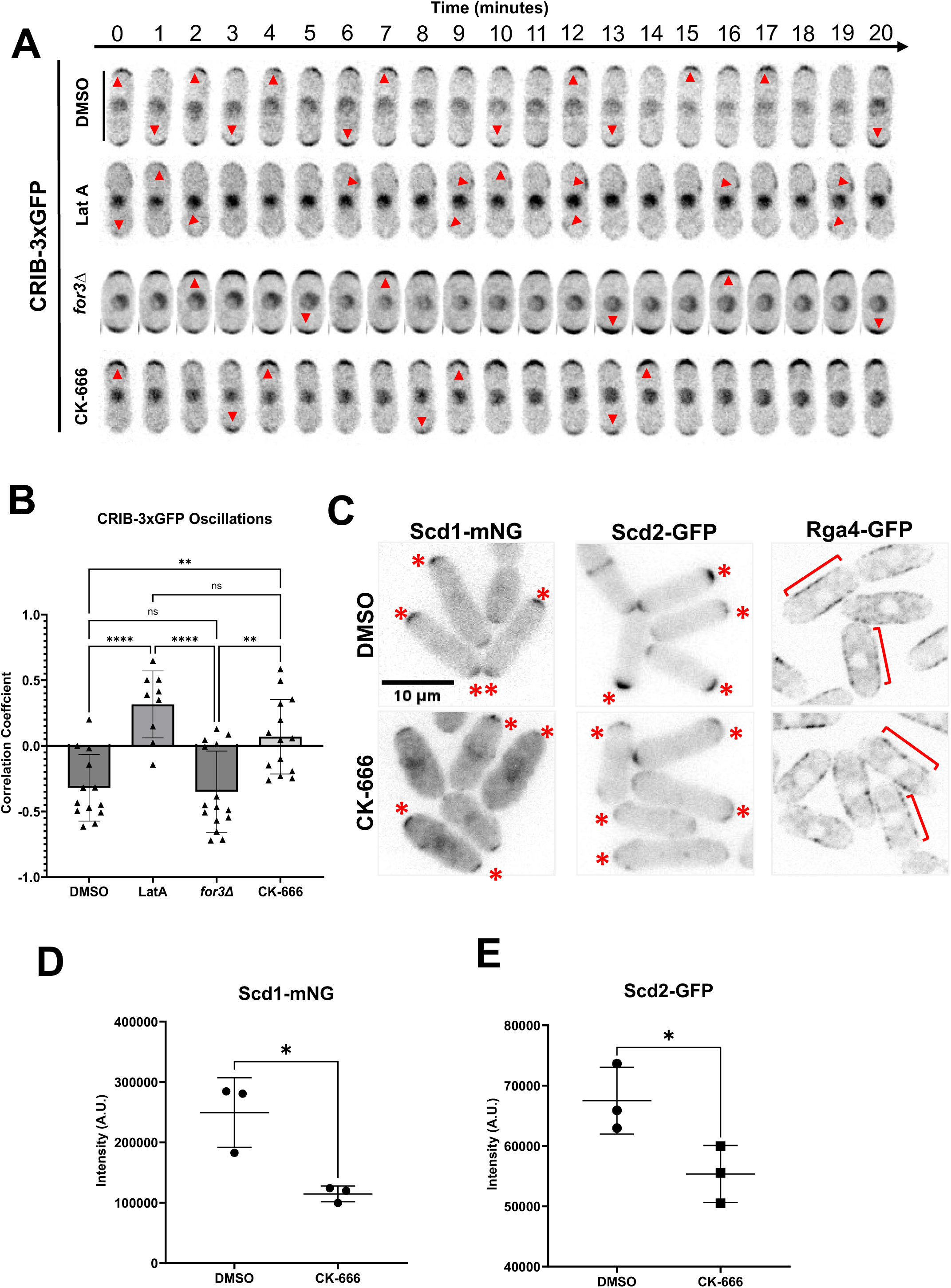
Branched actin is required for competition for active Cdc42 between growing ends. **A.** Cdc42 dynamics in CRIB-3xGFP expressing cells treated with DMSO, Lat-A, CK666 and in *for3Δ* mutants. Drug treatment and genetic perturbation were used to investigate the role of actin structures. Red arrow heads mark the site of Cdc42 activation. **B.** Correlation coefficient of active Cdc42 oscillations between cell ends. **C**. Localization of Scd1-mNG, Scd2-GFP and Rga4-GFP at the ends of DMSO and CK-666 treated cells. Red asterisks mark the site of Scd1-mNG and Scd2-GFP localization at the cell ends. Red brackets label Rga4-GFP along the cell sides. **D-E**. Quantification of Scd1-mNG and Scd2-GFP intensities at ends of DMSO and CK-666 treated cells. Scale bar, 10μm. n.s., not significant; p value, *<0.05, Student’s t-test.

Lat-A treatment disrupts all forms of F-actin structures including branched actin and linear actin cables [53, 56, 57]. Branched actin and actin cables are differentially nucleated and polymerized [40, 58–60]. Linear actin cables are nucleated by the formin, For3, whereas branched actin is nucleated by the Arp2/3 complex [40, 58–60]. The differential regulation between actin structures allows us to specifically target and deplete each structure for our investigations.

To this end, Cdc42 oscillations were analyzed in four conditions: DMSO, Lat-A, *for3Δ*, and CK-666. DMSO treatment served as a control (Fig.1A, B and S1A, B). *for3Δ* cells lack actin cables and cells treated with the Arp2/3 inhibiting drug CK-666 lack branched actin [40, 44, 59, 61, 62]. We then compared the level of anticorrelation between the cell ends in the absence of each actin structure (Fig.1B).

Interestingly, we find that *for3Δ* mutants still show anti-correlated active Cdc42 oscillations, similar to DMSO-treated cells, indicating that linear actin cables do not facilitate anticorrelation between the two ends (Fig.1B). However, cells treated with CK-666, which inhibits the Arp2/3 complex and branched actin formation, do not exhibit anti-correlated oscillations (Fig.1B). Moreover, in several cells treatment with CK-666 resulted in Cdc42 activation at mostly one end (Fig.1A and S1B). This suggests that the Arp2/3 complex promotes anticorrelation of active Cdc42 between the cell ends.

Previous works have demonstrated that under stress conditions, including Lat-A treatment, Cdc42 activity is depolarized and is instead localized along the cell sides [54, 55]. We asked if CK-666 treatment triggered a similar stress response resulting in the loss of anticorrelation. Stress response results in the activation of the mitogen-activated protein kinase (MAP kinase), Sty1 that promotes decreased localization of Scd1 at the cell ends and increased localization of the Cdc42 GEF Gef1 along the cell sides resulting in depolarized Cdc42 activation [19, 55]. Gef1 is mostly cytoplasmic and shows increased cortical localization only in response to stress [63]. We analyzed CRIB-3xGFP and Gef1-mNG localization in DMSO and CK-666 treated cells. In CK-666 treated cells, we did not observe CRIB-3xGFP localization along the cell sides (Fig.S2, A, B). Moreover, Gef1-mNG continued to display cytoplasmic localization in CK-666 treated cells (Fig.S2, C, D). This suggests that cells must lose all F-actin structures to trigger the stress response as opposed to just branched actin. To test this, we depleted branched actin in mutants lacking linear actin cables to mimic Lat-A treatment. We treated *for3Δ* mutants with CK-666 to get rid of all F-actin structures. Indeed, we find that *for3*Δ cells when treated with CK-666 displayed depolarized CRIB-3xGFP and Gef1-mNG (Fig.S2). This suggests that CK-666 treatment by itself does not trigger a stress response.

### Inhibition of the Arp2/3 complex depletes Scd1 from the cell ends

Next, we elucidated how inhibition of the Arp2/3 complex disrupted Cdc42’s regulation resulting in loss of anticorrelation between the cell ends. The Arp2/3 complex is required for branched actin-mediated endocytosis and CK-666 treatment is known to abolish endocytosis [42, 47, 48, 62, 64]. We asked if the loss of endocytosis disrupted Cdc42 regulators resulting in loss of bipolarity and anticorrelation. Endocytosis is required to recycle proteins from the cell cortex [65, 66]. Inhibition of endocytosis leads to accumulation of these recycled proteins at the cell cortex. We asked if the Cdc42 regulators were similarly recycled by endocytosis. We visualized fluorescently labeled Cdc42 regulators in DMSO treated control cells and cells treated with CK-666 (Fig.1C). Scd1-mNG (primary GEF), Scd2-GFP (Scd1 Scaffold), and Rga4-GFP (Primary GAP) were imaged using fluorescent microscopy (Fig.1C) [20, 23]. The Cdc42 GAP, Rga4-GFP did not show any change in its localization in CK-666 treated cells. Contrary to the expectation of endocytosis-driven recycling, we find that Scd1, and its scaffold Scd2, are significantly depleted from growing ends in cells upon Arp2/3 complex inhibition (Fig.1D, E). Since these regulators do not accumulate at the cell cortex upon CK-666 treatment, it suggests that they are not recycled from the cell cortex by endocytosis. Rather, endocytosis appears to enhance localization of Scd1 and Scd2 at the cell ends. Scd1 levels are known to oscillate between the cell ends similar to active Cdc42 [11]. As a result, when imaging Scd1 at any given point of time, it either accumulates at any one cell end or is distributed between the two ends. Indeed, in a DMSO treated population, snapshots of Scd1-mNG show 53% of cells with bipolar localization. In CK-666 treated cells, in addition to decrease in levels of Scd1-mNG, we also observe bipolar localization in only 37% of cells. This suggests that in the absence of branched actin, Scd1 localization is restricted possibly due to the disruption of its oscillation.

### Cdc42-GTP inhibits the accumulation of Scd1 through an intermediary molecule

To relate our observations on polarization regulators under perturbed conditions to the core oscillator controlling Cdc42-GTP dynamics in a rigorous manner, we constructed two models to explore potential mechanisms at the systems level. With a framework combining ordinary and partial differential equations (ODE-PDE), a previous model used an assumption of Cdc42 auto-catalysis and an additional negative feedback loop between Cdc42 and Scd1 to explain Cdc42 oscillation (a classical structure chemical oscillator) [67, 68]. However, there is no experimental evidence supporting the auto-catalytic function of Cdc42 to date. We therefore removed this assumption and implemented a regulatory network (Model 1) describing Scd1-mediated Cdc42 activity cycle, an experimentally supported positive regulation of Scd1 by Cdc42-GTP, and a hypothetical, direct negative feedback loop between Cdc42-GTP and Scd1 (Negative feedback loop is required for oscillation) (Fig.2A.i.). In this ODE-PDE model, the two ends of the cell contain reactions at the membrane (Fig.2A), whereas the molecules in the cytoplasm or at the membrane on the sides are only allowed to diffuse [68]. The ODE-PDE model therefore has three pools of Cdc42 and its regulators, one at each cell end and the third in the cytoplasm. We assign fixed concentrations of Cdc42 and the GEF Scd1, distributed between the three pools depending on their recruitment and removal rates (Fig.3B). With 100,000 randomly sampled parameter values, this model failed to produce oscillatory dynamics observed experimentally (Fig.2A i, Model 1) (see Methods). This suggests that there exists another regulator essential for oscillation. It was known that delayed negative feedback can produce oscillations without auto-catalysis [69]. Furthermore, membrane bound Scd1 inhibitors have been found experimentally [12]. We therefore introduced an additional unknown protein X to our model (Model 2). Protein X is activated/recruited by active Cdc42, and it inhibits the accumulation of Scd1 at the membrane, which gives rise to a delayed feedback loop. In contrast to Model 1, we found that 1.4% of randomly sampled parameter sets of Model 2 produced oscillation (Fig.2A.iii) (see Methods). Since Model 2 captured our experimental observations with respect to Cdc42 oscillatory dynamics, we used this model and a representative oscillation-enabling parameter set to further investigate the role of branched actin in establishing anticorrelation. We assumed that the removal rate constants (δ) of each molecule positively correlate with endocytosis because branched actin in fission yeast is required for endocytosis. The scenario of branched actin disruption is described by reduced removal rate constants in our model (i.e., decreased δ) (Fig.2 C.i). To explore the influence of branched actin, we adjusted the removal rate constants of Scd1, Cdc42, and protein X at the ends individually with the representative parameter set in Model 2 (Fig.2C). We found that decreasing the removal rates of Cdc42 dampened the overall signal of Cdc42 as well as the GEF Scd1 and factor X (Fig.2C iii left), but the steady state distributions of these molecules are symmetrical at the two ends. This bipolar distribution was also observed when Scd1 removal rate was decreased (Fig.2C, iii middle). The bipolarity was observed in a wide range of removal rate constants (Fig.S3A-B). These simulations did not match our experimental observations of monopolar distribution upon loss of branched actin (Fig.1). In contrast, decreasing the removal rate constant of factor X showed monopolar dynamics of Cdc42, Scd1 and factor X itself (Fig.2C iii right). Furthermore, we found that the mean level of Scd1 decreased with this perturbation, a phenomenon qualitatively reflected in our experiment (black dashed line in Fig.2C iii, and Fig.1C and D). Overall, our simulation results showed that removal of protein X (i.e., the Scd1 inhibitor) is important for maintaining cell bipolarity and Cdc42 oscillations at both ends.

**Figure 2.**
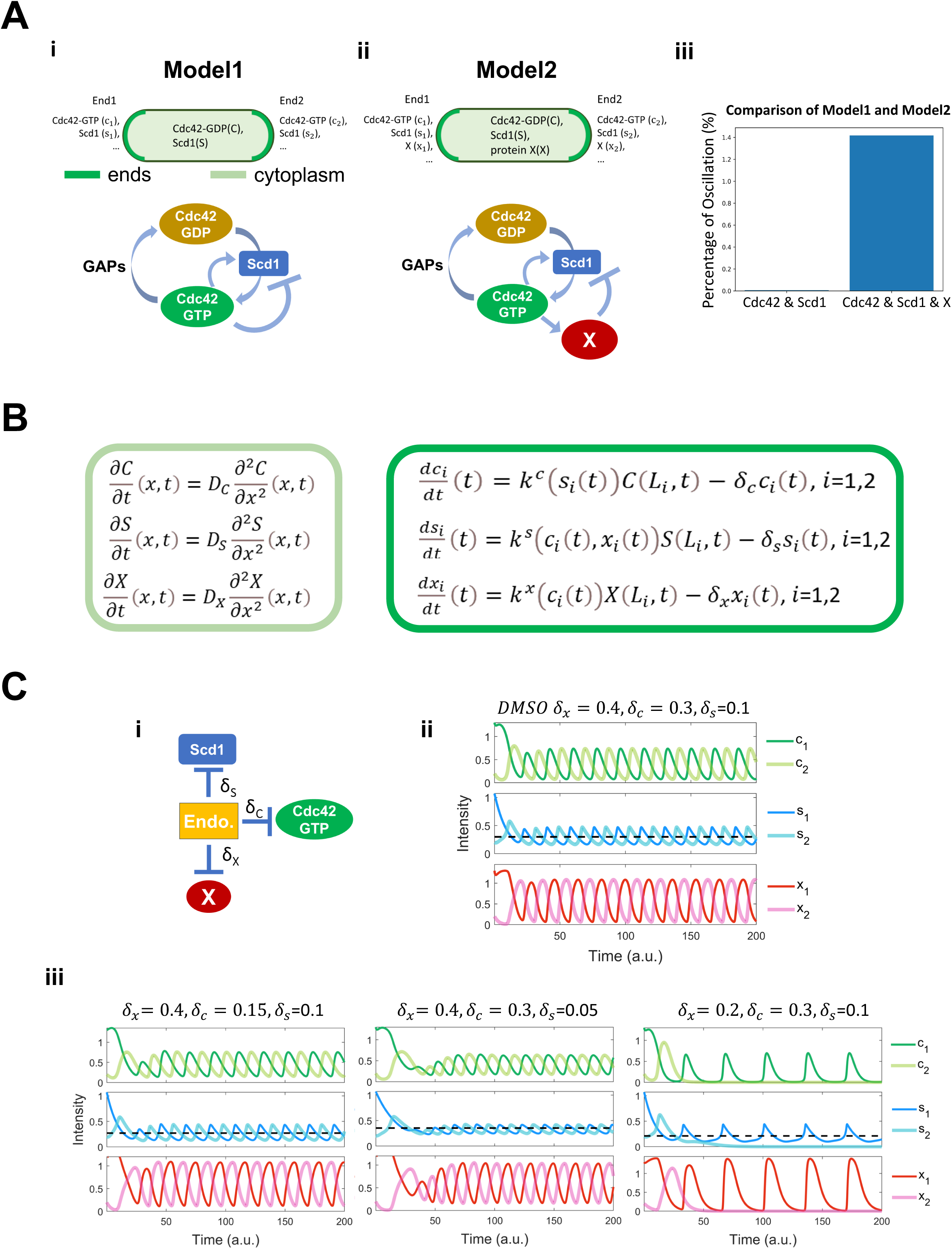
Models for bipolar and monopolar dynamics involving endocytosis. **A.** (i) Membranes at the cell ends accumulate Cdc42-GTP and Scd1, while the cytoplasm contains diffused Cdc42-GDP and Scd1. The panel below describes the Cdc42 activation network corresponding to Model1. Model1 incorporates feedback loops involving Cdc42-GTP, Cdc42-GDP, and GEF (Scd1), whereby the presence of Cdc42-GTP directly inhibits the membrane accumulation of GEF. (ii) Membranes at the cell ends accumulate Cdc42-GTP, Scd1, and an unknow protein X, while the cytoplasm contains diffused Cdc42-GDP, Scd1 and X. The panel below describes the Cdc42 activation network corresponding to Model2. Model2 incorporates feedback loops involving Cdc42-GTP, Cdc42-GDP, GEF (Scd1), and an unknown protein X, whereby the presence of Cdc42-GTP inhibits the membrane accumulation of GEF via protein X. (iii) A comparison between the percentage of oscillatory dynamics in Model1 and Model2. The number of parameter sets for 2 models is 100,000. **B.** Reaction-diffusion equations for Model2 (details in supplemental materials). Light green frame represents protein diffusion in cytoplasm, and dark green frame denotes protein production and degradation at the cell ends. **C.** (i) The effects of endocytosis on each protein in the Cdc42 activation network. (ii) Numerical results of the PDE-ODE simulations for Model 2. (iii) Numerical results for Model 2 were obtained by varying the removal rates of each protein. The black dashed line is the average concentration of Scd1 at the oscillatory dynamics tip. Initial conditions at each cell end 1 and 2: c1=1.3, c2=0.2, s1=1.07, s2=0.2, x1=1.3, x2=0.2 (c=Cdc42, s=Scd1, x= protein X).

**Figure 3.**
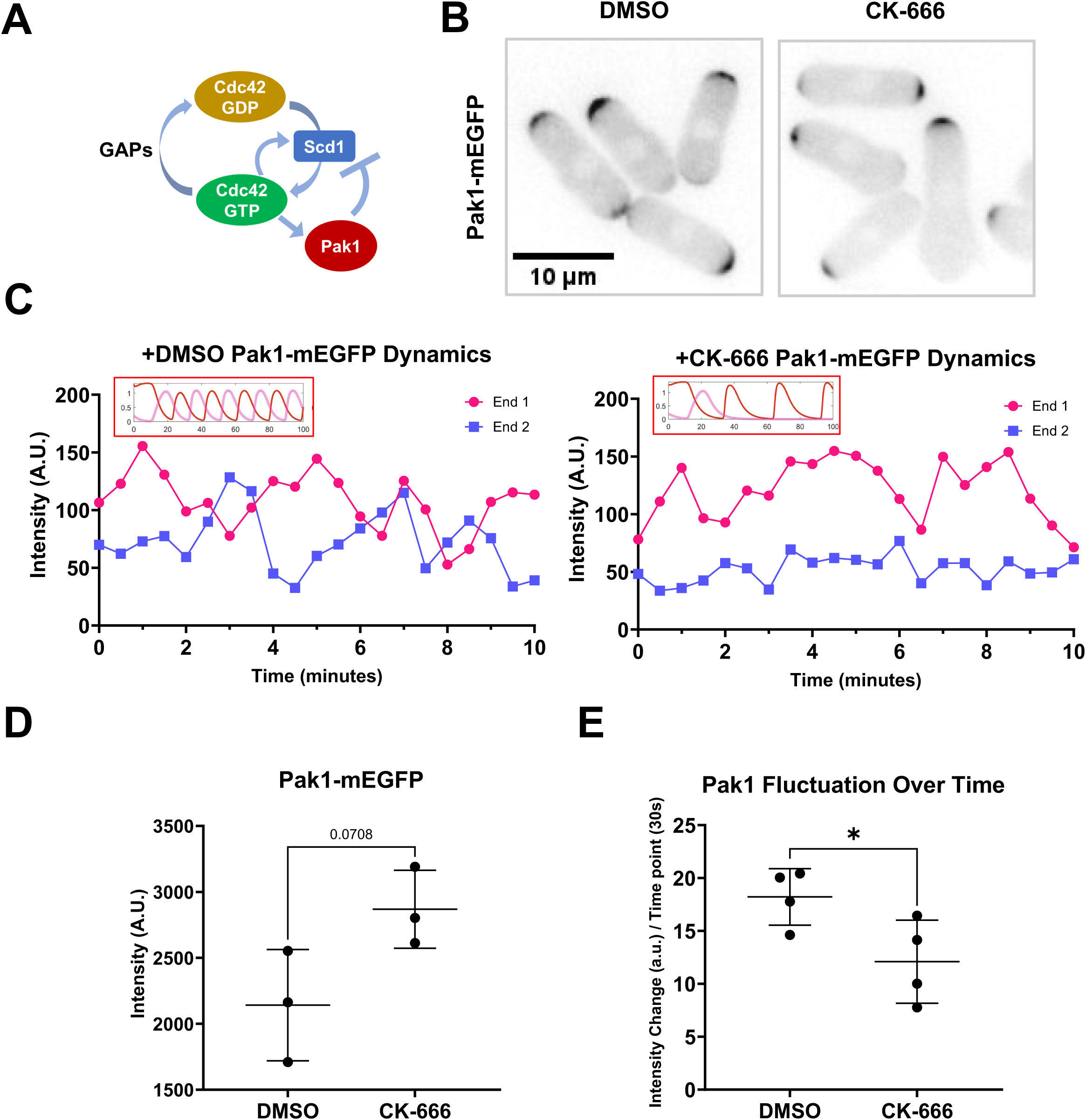
The Pak1 kinase, a potent inhibitor of Scd1, becomes more stable when branched actin is blocked. **A.** Model describing Cdc42 activation network where Pak1 kinase upon activation by Cdc42 triggers Scd1 removal. **B.** Pak1-mEGFP localization in DMSO and CK-666 treated cells. **C.** Pak1-mEGFP dynamics in cells with treated with DMSO or CK-666. Dynamics observed *in vivo* resembled the dynamics seen in model predictions (insets) **D.** Quantification of Pak1-mEGFP localization at the brightest cell end in DMSO or CK-666 treated cells. **E.** Quantification of the extent of Pak1-mEGFP fluctuations at the cell ends. Scale bar, 10µm. p-value, *<0.05, Student’s t-test.

### The PAK kinase, Pak1, stabilizes when the Arp2/3 complex is inhibited

Our model predicts that decreasing the removal of the inhibitor X leads to asymmetric Cdc42 dynamics and decreased Scd1 at the cell ends. It has been reported that the Cdc42 effector kinase Pak1 is part of the time-delayed negative feedback and prevents Scd1 localization [28–30]. Thus, factor X could be Pak1 kinase in Model 2. Pak1 is an essential protein and is involved in regulating endocytosis via phosphorylation of the type 1 Myosin, Myo1 [70]. Pak1 localization to the cell ends is dependent on active Cdc42 (Fig.3A and Fig.S4). Cells with decreased active Cdc42 due to *scd1Δ* show a decrease in Pak1-mEGFP at the cell cortex (Fig.S4). In *gef1Δ* mutants Pak1-mEGFP is mostly monopolar, while in the *rga4Δrga6Δ* mutant Pak1-mEGFP at the cell cortex is enhanced (Fig.S4). Similar to active Cdc42 dynamics, Pak1-mEGFP displays anticorrelated oscillations between the two cells ends (Fig.3B). We posit that when activated by Cdc42, Pak1 promotes endocytosis and this leads to its own displacement from the cell ends. Thus, we predict that in the absence of branched actin, Pak1 levels at the cell ends will stabilize and decrease in fluctuation. To test this, we used Pak1-mEGFP to visualize Pak1 localization and dynamics using timelapse microscopy. We find that upon CK-666 treatment, Pak1 dynamics at the cell ends resemble that of our model 2 with decreased removal rate of factor X (Fig.3B and Fig.2Ciii). We next measured the intensity of Pak1-mEGFP at the cell ends in DMSO and CK-666 treated cells. In snapshot images we did not see a significant increase in Pak1-mEGFP levels upon CK-666 treatment, although we saw a trend in that direction (Fig.3C). However, when we analyzed Pak1-mEGFP dynamics at the cell ends over time we observed that it stabilizes with decreased fluctuations in CK-666 treated cells (Fig.3B, C, D and S6A).

Our model predicts that stabilizing Pak1 kinase leads to decreased Scd1 levels at the cell ends. To test this, we investigated if Scd1 intensity would be impacted upon CK-666 treatment in the absence of Pak1 activity. To this end, we measured Scd1-mNG intensities in the hypomorphic *pak1* mutant *orb2-34* (*pak1-ts*) (Fig.4A). The *pak1-ts* mutant cells are monopolar and wider even at the permissive temperature (25°C). Moreover, in keeping with previous reports, Scd1-mNG levels at the cell ends increases in *pak1-ts* mutant at permissive temperature (Fig.4A, B). We treated *pak1-ts* cells at permissive temperature with DMSO and CK-666 and compared Scd1-mNG intensity in these cells against *pak1+* controls under the same conditions. We find that Scd1-mNG intensity is not depleted in *pak1-ts* cells treated with CK-666 (Fig.5A, B). This suggests that the decrease in Scd1 levels observed in CK-666 treated cells is indeed due to Pak1 kinase activity. The *pak1-ts* cells are characteristically monopolar, however we unexpectedly find that around 40% of these cells treated with CK-666 showed bipolar localization of Scd1-mNG (Fig.4A, asterisks). The mechanism of this bipolar phenotype in the absence of Pak1 is not yet understood with our existing model: in Model 2, Pak1 (X in Fig.2) mediated negative feedback is essential for bipolarity (Fig.S5). We therefore included an additional, hypothetical negative feedback loop involving Cdc42 in our model (Model 3) (Fig.4Ci) (see details in supplementary materials). Figure 4Cii showed the simulation results of Model 3 under both DSMO and CK666 conditions. As expected, the modeling shows monopolar dynamics in the absence of Pak1. However, the simulations captured both bipolar and monopolar phenotypes, when removal rates of Cdc42 and Scd1 are reduced, suggesting a heterogeneous response to CK-666 in the absence of Pak1 (Fig.4D).

**Figure 4.**
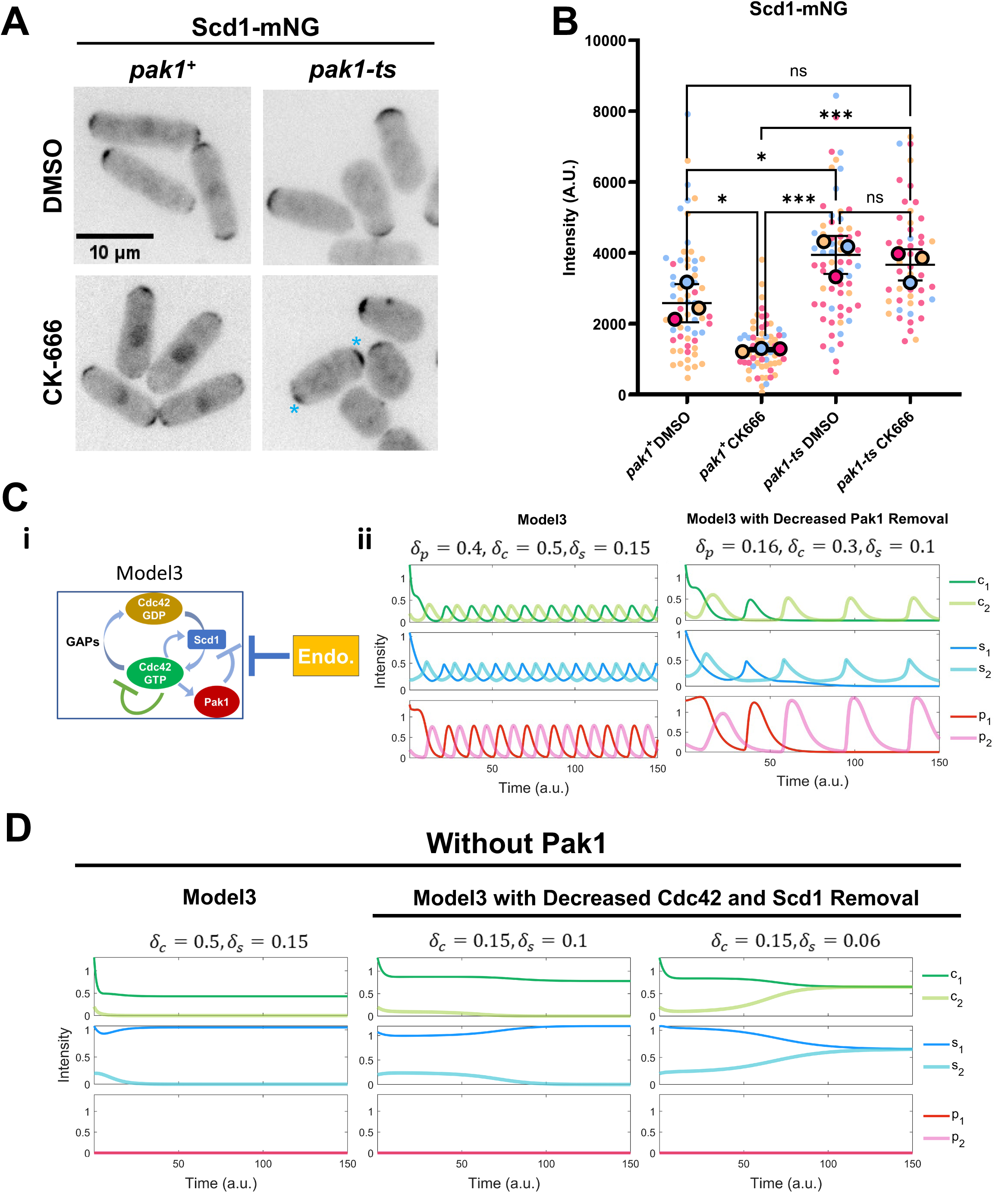
Loss of proper Scd1 localization is caused by Pak1 stabilizing. **A.** Scd1-mNG localization in *pak1+* and *pak1-ts* mutant cells. Blue asterisks depict bipolar Scd1-mNG localization. **B.** Quantification of Scd1-mNG intensities at the cell ends in *pak1+* and *pak1-ts* cells treated with DMSO and CK-666. Colors represent each replicate by day. Large circles are the means of each experimental replicate (n ≥ 10 cells). **C.** (i) Model3 incorporates feedback loops involving Cdc42-GTP, Cdc42-GDP, GEF (Scd1), Pak1, and endocytosis, and an additional negative signaling pathway inhibiting Cdc42-GTP. (ii) Left panel shows Model3 simulations, and the right panel shows simulations with decreased removal rate of Pak1. **D.** Model3 simulations with an additional negative signaling pathway simulated via decreased removal rate of Cdc42 and Scd1. Scale bar, 10µm. p-value, *<0.05, ***<0.0003, One way ANOVA, followed by Tukey’s multiple comparison test.

**Figure 5.**
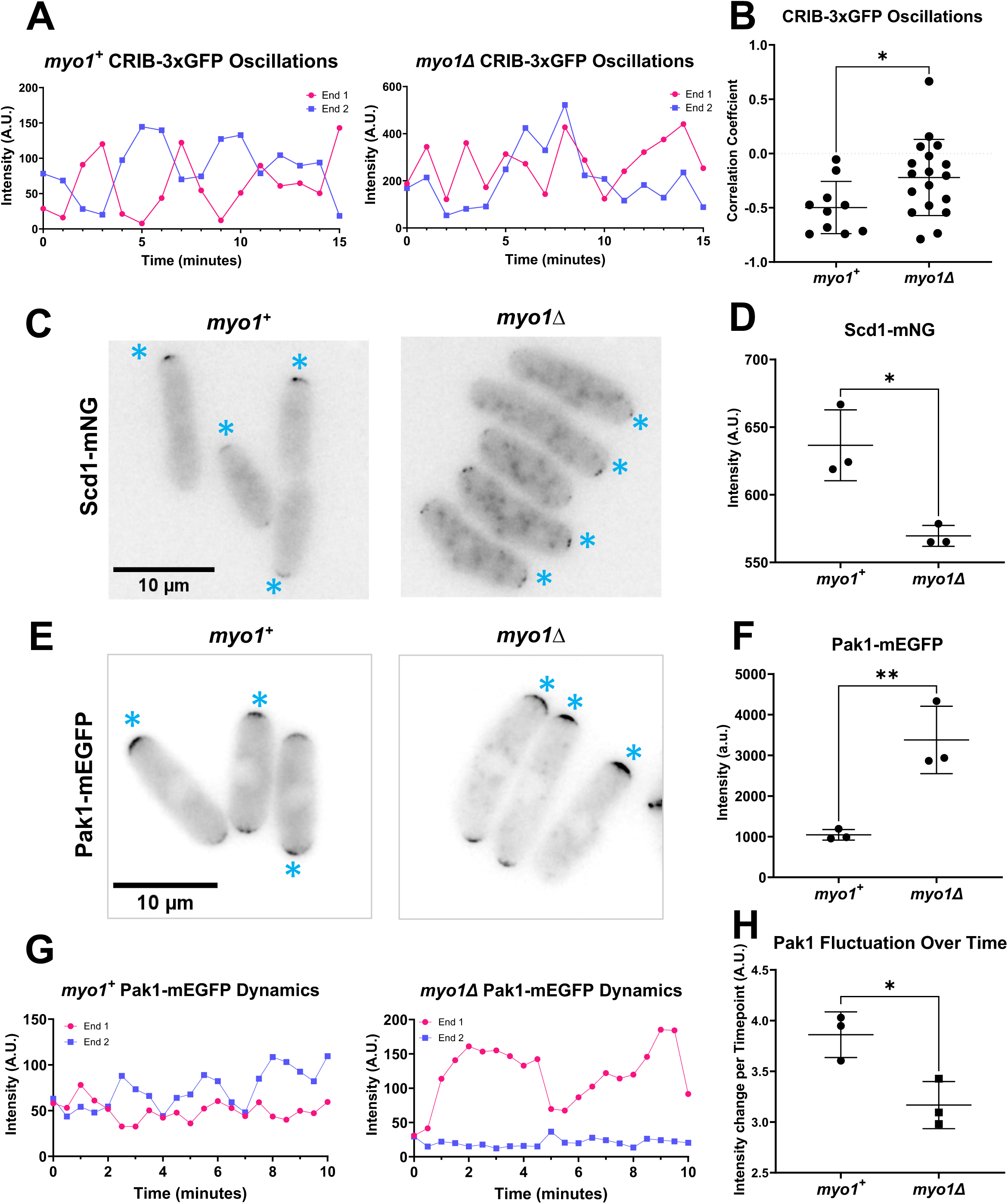
Endocytic mutant *myo1Δ* cells show disrupted Cdc42 activation dynamics similar to CK-666 treatment. **A.** CRIB-3xGFP dynamics were observed in *myo1+* and *myo1Δ* cells. **B**. Correlation coefficients of active Cdc42 oscillations between cell ends in *myo1+* cells and *myo1Δ* mutants. **C**. Scd1-mNG localization in *myo1+* and *myo1Δ* cells. Asterisks indicate Scd1-mNG localization at the brighter cell end. **D**. Quantification of Scd1-mNG accumulation at cell ends in *myo1+* and *myo1Δ* cells. **E.** Pak1-mEGFP localization in *myo1+* and *myo1Δ* cells. Asterisks indicate Pak1-mEGFP localization at the brighter cell end **F.** Quantification of Pak1-mEGFP accumulation at cell ends in *myo1+* cells and *myo1Δ* mutants. **G.** Pak1-mEGFP dynamics at cell ends in *myo1+* and *myo1Δ* cells. **H.** Quantification of the extent of Pak1-mEGFP fluctuation in *myo1+* and *myo1Δ* cells. Scale bar 10µm. p-value, *<0.05, **<0.0085, Student’s t-test.

### Loss of the Arp2/3 regulator *myo1* disrupts Cdc42 oscillatory dynamics

Our observations show disruption of the Cdc42 oscillatory dynamics in CK-666 treated cells. CK-666 is an inhibitor of Arp3 and the Arp2/3 complex is required for endocytosis. Thus, we posit that endocytosis is required for maintaining proper Cdc42 oscillatory dynamics. If true, then we should also see similar disruption of Cdc42 oscillatory patterns in endocytic mutants. Endocytosis is an essential cellular process and most endocytic mutants show severe cell growth and polarity defects thus complicating such an analysis. The Arp2/3 complex regulator *myo1* is not essential and loss of *myo1* results in shape and growth defects but these cells are still able to polarize [47]. Thus, we used the *myo1Δ* mutant to investigate Cdc42 oscillatory dynamics. We find that *myo1Δ* mutants show similar defects in Cdc42 oscillatory dynamics to those observed in CK-666 treated cells (Fig.5A, B). We observed Cdc42 oscillations in *myo1+* and *myo1Δ* mutants expressing CRIB-3xGFP. Normally, cells show anticorrelated CRIB-3xGFP oscillations with a correlation coefficient of about −0.5 (Fig.5B). In *myo1Δ* cells CRIB-3xGFP signals at the cell ends continued to fluctuate but displayed an average correlation coefficient of about −0.2 indicating a decrease in anticorrelation (Fig.5B and S6B). Next, we compared the localization of Scd1-mNG at the cell ends in *myo1+* and *myo1Δ* cells. Similar to CK-666 treated cells, *myo1Δ* mutants showed a decrease in Scd1-mNG levels at the cell ends (Fig.5C, D). We also observed increased monopolar localization of Scd1-mNG at the cell ends in *myo1Δ* mutants. Our results with CK-666 treated cells suggest that loss of Scd1 levels at the cell ends is due to the stabilization of Pak1 kinase at those ends. Thus, we analyzed Pak1-mEGFP levels and dynamics at the cell ends in *myo1Δ* mutants. We find that in *myo1Δ* mutants Pak1-mEGFP levels at the cell ends increase (Fig.5E, F). We observed Pak1-mEGFP dynamics using timelapse imaging in *myo1+* and *myo1Δ* cells. Pak1-mEGFP localization mostly appeared at one cell end and displayed decreased fluctuations over time in *myo1Δ* cells (Fig.5G, H and S6B). This suggests that, similar to CK-666 treated cells, Pak1 dynamics at the cell ends stabilized in *myo1Δ* cells. Together, these data suggest that efficient endocytosis is required for proper Cdc42 activity between the cell ends.

### Endocytic events deplete Pak1 from the membrane

Our data suggests that the Arp2/3 complex is required for Pak1 removal to allow Scd1 localization and anticorrelated oscillations for Cdc42 between the cell ends. As a result, we hypothesize that Pak1 is depleted from the cell end via endocytosis. Pak1 kinase has been shown to phosphorylate Myo1 during endocytosis. However, Pak1-mEGFP does not localize exclusively to the endocytic patches, but rather appears all over the cell end. This is in agreement with the fact that Pak1 localization at the cell ends is dependent on active Cdc42 and Pak1 regulates other pathways in addition to endocytosis [30, 71]. Thus, we posit that only Pak1 kinase molecules overlapping with the endocytic patches are removed via endocytosis. To test this, we studied the relationship between endocytic events and the loss of Pak1 from the membrane.

Endocytic patches in fission yeast can be detected using the actin crosslinking protein Fimbrin, Fim1 [72]. Fim1 specifically localizes to branched actin patches and internalizes into the cytoplasm with the endocytic vesicle during endocytosis [72]. We analyzed Fim1-mCherry internalization in cells also expressing Pak1-mEGFP. We observe that while the Fim1-mCherry labeled endocytic patch internalizes into the cytoplasm, Pak1-mEGFP does not (Fig.6A). However, we observed Pak1-mEGFP overlapping with Fim1-mCherry is lost from the plasma membrane when the endocytic patch internalizes. Most endocytic proteins are known to internalize with the vesicle. However, several regulators of endocytosis do not internalize but are simply lost from the membrane when the vesicle internalizes. In fission yeast, the endocytic proteins such as the F-BAR containing Cdc15 and Myo1 are lost from the membrane upon vesicle internalization [73, 74]. In mutants with a delay in vesicle internalization, loss of these proteins from the plasma membrane is also delayed [42]. It is possible that Pak1 also shows a similar loss at the membrane during vesicle internalization. To test this, we measured the intensity of Fim1-mCherry and Pak1-mEGFP simultaneously at the plasma membrane. We observe that each time Fim1-mCherry starts to internalize (Fig.6C, dashed lines), Pak1-mEGFP is also lost from the plasma membrane as indicated by a decrease in its intensity at the membrane (Fig.6A, B and C). Next, we quantified when Pak1-mEGFP was lost from the membrane compared to Fim1-mCherry internalization (Fig.6C, D). We find that, on average, Pak1-mEGFP is lost from the membrane within 1 second of Fim1-mCherry internalization. This suggests that Pak1 is removed from cell ends when the endocytic vesicle internalizes.

**Figure 6.**
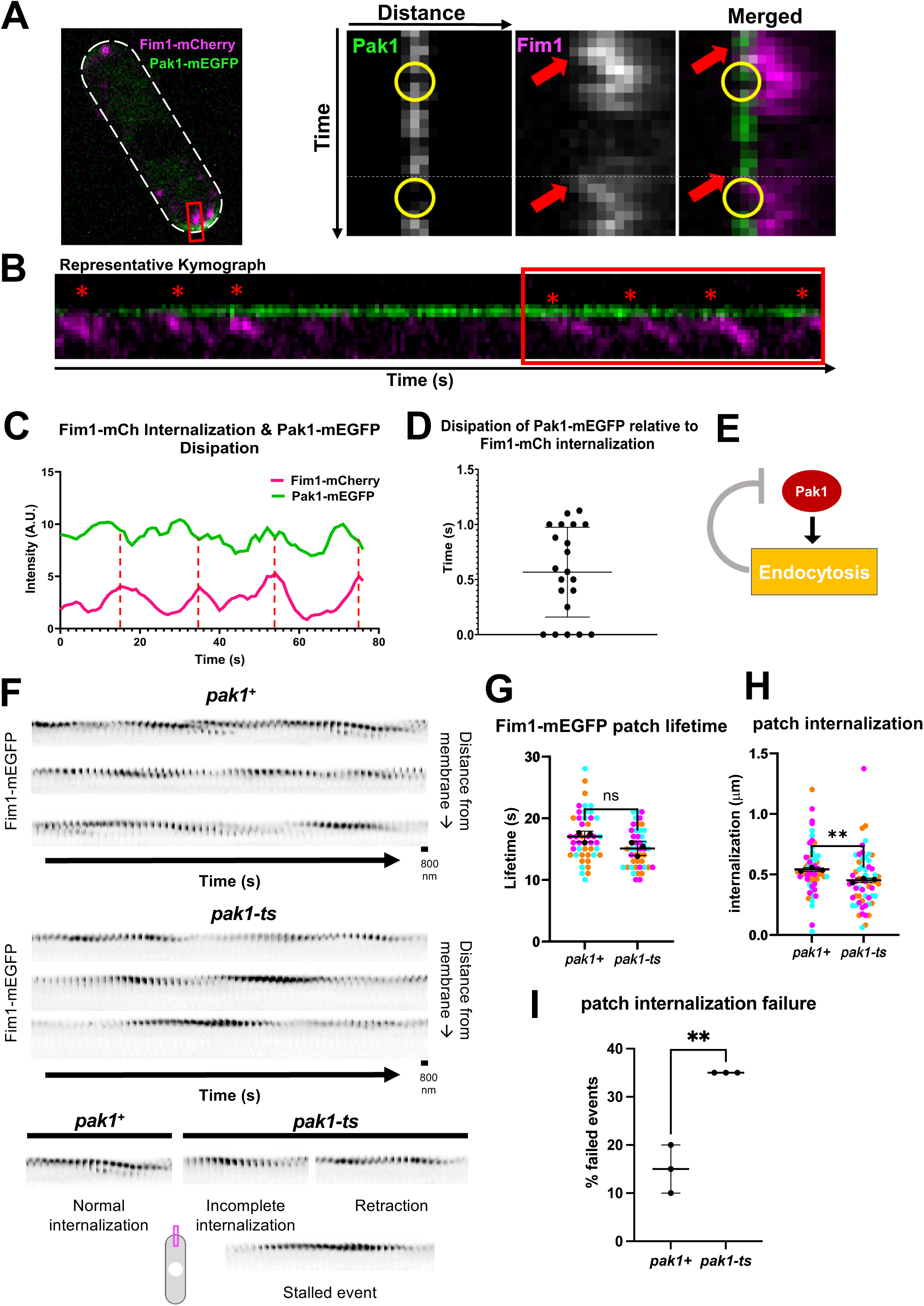
Pak1 is removed from the cell end via endocytosis. **A.** Fim1-mCherry and Pak1-mEGFP at cell ends during endocytic events. Left panel shows the whole cell with a white dashed outline. Red boxes mark the region shown as kymographs in the right panels. Yellow circles mark the Pak1-mEGFP loss from the membrane while red arrow marks the onset of Fim1-mCherry internalization at the membrane. **B.** Kymograph of Pak1-mEGFP and Fim1-mCherry as captured in a 3-minute timelapse movies with 1 second intervals. Red box marks the region quantified in C. **C.** Quantification of Pak1-mEGFP and Fim1-mCherry intensities at the membrane over time. Red dashed lines indicate the peak and subsequent drop of Fim1-mCherry intensity. **D.** Quantification of the time taken for dissipation of Pak1-mEGFP relative to Fim1-mCherry internalization. (n ≥6 endocytic events per kymograph, N= 21 kymographs from 3 experiments) **E.** Schematic depicting our hypothesis that Pak1 activity promotes its own removal through endocytosis. **F.** Montages of Fim1-mEGFP labeled endocytic patches at growing ends of interphase *pak1^+^* and *pak1-ts* cells (scale bars = 800 nm; frame rates = 1 second per frame (SPF)). **G.** Quantification for lifetimes of Fim1-mEGFP labeled endocytic patches in *pak1^+^* and *pak1-ts* (n=15 endocytic patches per experiment, N=3 experiments). **H.** Quantification of Fim1-mEGFP labeled endocytic patch internalization into the cell interior from the plasma membrane in *pak1-ts* mutants compared to *pak1^+^* cells (n=20 endocytic patches per genotype per experiment, N=3 experiments). **I.** Quantification of failed endocytic events in *pak1^+^* cells compared to *pak1-ts.* Failed events do not internalize beyond 350 nm from the plasma membrane (n=20 endocytic patches per genotype per experiment, N=3 experiments). Symbol colors in graphs = distinct experiments. Solid symbols = means of experiments. n.s., not significant; p-value, **<0.01, Student’s t-test.

### Pak1 activity promotes successful internalization of endocytic vesicles

Our model indicates that Pak1 removal is dependent on its accumulation at the cell ends. This suggests that Pak1 activity drives endocytosis which promotes its own removal. While the role for Pak1 kinase in endocytosis has not been fully investigated, it has been shown to phosphorylate Myo1 during endocytosis [47, 70]. Thus, we expected that Pak1 kinase is required for endocytosis. To test this, we analyzed endocytic dynamics in the absence of Pak1 kinase. The *pak1-ts* mutant, *orb2-34* shows polarity defects even at its permissive temperature of 25°C. We find that successful endocytic events are significantly reduced in *pak1-ts* mutants (Fig.6F, I) at 25°C. Fim1-mEGFP was used to specifically visualize endocytic patches. We observed endocytic vesicles still form at the membrane in *pak1+* and *pak1-ts* cells and their lifetime at the cell membrane remains the same (Fig.6F, G). Thus, Pak1 does not influence the ability of endocytic vesicles to form. Endocytic vesicles that internalize >350 nm away from the membrane are considered to have undergone successful scission from the plasma membrane [75]. We find that there is a significant decrease in the overall distance that vesicles internalize in *pak1-ts* mutants compared to *pak1+* cells (Fig.6F, H). Furthermore, a higher fraction of the patches in *pak1-ts* mutants did not internalize beyond 350nm indicating a greater fraction of failed endocytic events compared to *pak1+* cells (Fig.6F, I). In addition, vesicles that do not properly internalize show three types of failed endocytic events such as incomplete internalizations, vesicle retractions, or stalled events (Fig.6F).

### Pak1 accumulation at the cell ends lags behind Cdc42 activation

Oscillatory patterning requires positive feedback and time-delayed negative feedback [76]. Cdc42 positive feedback comprises of Scd1 and Scd2 while negative feedback is mediated by the Pak1 kinase [18, 21, 77, 78]. To this end, our model predicts that there is a spatiotemporal delay (phase shift) between the recruitment and peak accumulation of Cdc42 positive feedback proteins and the negative feedback protein, Pak1 at the growing cell ends (Fig.7A). Indeed, advancements in live cell microscopy have allowed for rapid timelapse imaging of even sparse proteins such as Scd1 *in vivo*.

**Figure 7.**
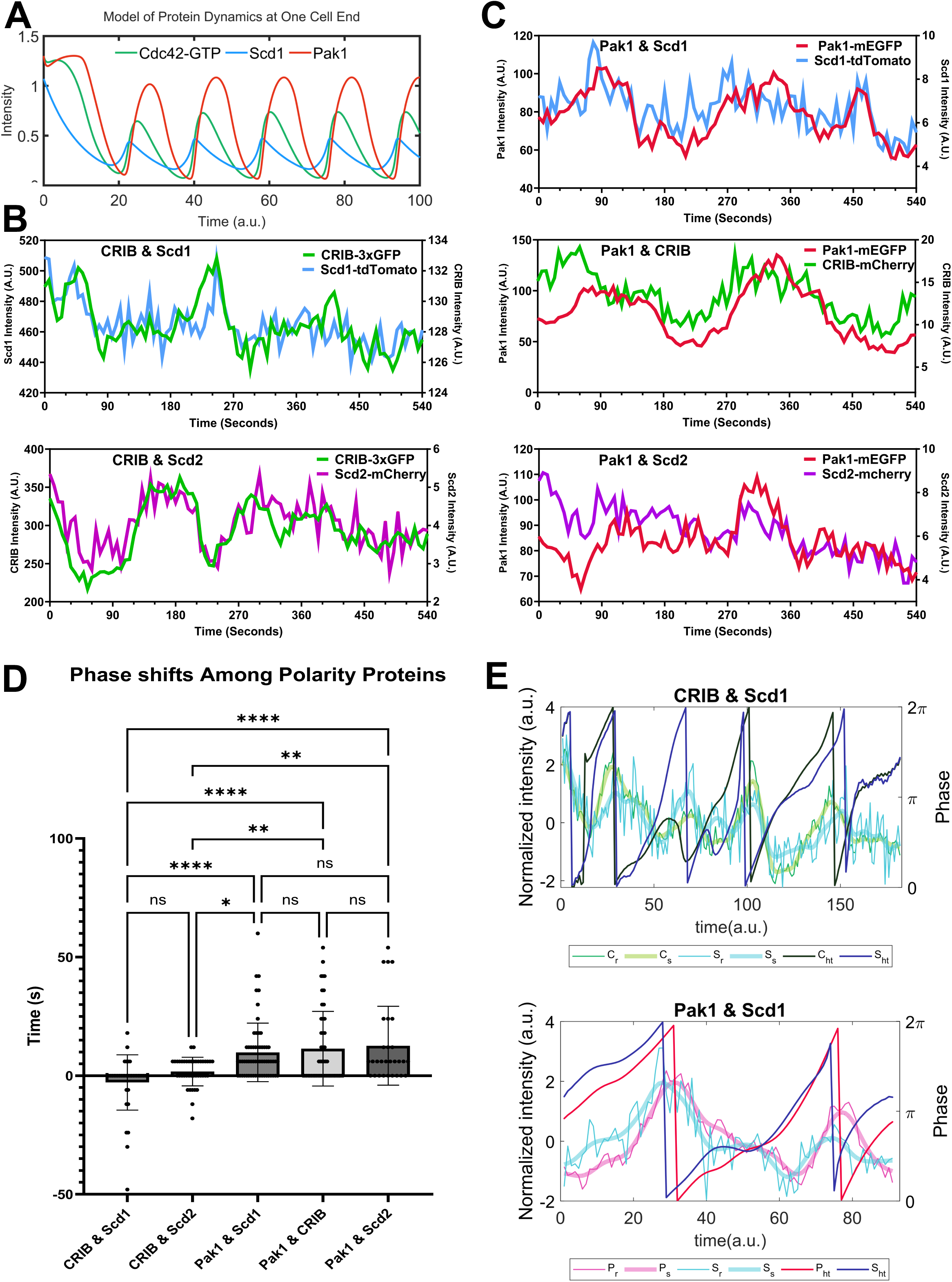
There is an observable phase shift between positive polarity proteins and Pak1 at the cell end. **A.** Simulations of our models show a delay or phase shift between the peak accumulations of Cdc42-GTP and Pak1 at the growing end. **B.** Pairs of positive-feedback polarity proteins (CRIB-3xGFP and Scd1-tdTomato, CRIB-3xGFP and Scd2-mCherry) were simultaneously observed at the cell end. **C.** Each positive-feedback protein (Scd1-tdTomato, CRIB-mCherry, and Scd2-mCherry) was observed simultaneously with the negative-feedback protein, Pak1-mEGFP. **D.** The shift in time required to get the best correlation coefficient between signals from each pair of proteins was quantified using cross-correlation analysis. **E.** The Hilbert transform applied to the smoothened traces of Cdc42-GTP and Scd1 for their phase reconstruction and phase shift (Top). The Hilbert transform applied to the smoothened traces of Pak1 and Scd1 for their phase reconstruction and phase shift (Bottom). The thinnest traces represent normalized raw data, the thickest traces represent smoothened traces based on the normalized data, the darkest traces represent the phase progression achieved through Hilbert transform. n.s., not significant; p-value, **<0.005, ****<0.0001, One way ANOVA, followed by Tukey’s multiple comparison test.

To test this, we used strains with a combination of fluorescently tagged CRIB-3xGFP or CRIB-mCherry, Scd1-mNG or Scd1-tdTomato, Scd2-mCherry, and Pak1-mEGFP (Fig.7B, C). We find that there is practically no delay (average of 2 seconds) between positive feedback proteins such as CRIB-3xGFP & Scd1-tdTomato and CRIB-3xGFP & Scd2-mCherry (Fig.7B, D and S7A). However, there is an average delay of 12 seconds between the accumulation of positive feedback proteins (CRIB-mCherry, Scd1-tdTomato, or Scd2-mCherry) and the peak accumulation of Pak1-mEGFP (Fig.7C, D and S7B). We validated these results *in silico* (Fig.7E). The phase shift diagrams were generated using the Hilbert Transform of the normalized data of proteins at the ends. These analyses show the raw traces and transformed traces of the protein levels. The latter highlights the oscillatory cycle of pairs of proteins and assesses for any existing phase shifts (Fig.7D) [79]. As we expected the GEF Scd1-tdTomato peaks are in phase with those of active Cdc42 labeled by CRIB-3xGFP. Further, Pak1-mEGFP accumulation had a phase shift with respect to Scd1-tdTomato accumulation.

## DISCUSSION

While Cdc42 activation spatiotemporally regulates actin organization for membrane trafficking and polarized growth, it is not well understood how Cdc42 itself is dynamically regulated. In fission yeast, Cdc42 and its regulators undergo oscillatory dynamics between the two cell ends, and this lends itself to dynamic regulation of actin organization [13, 17, 46, 61, 80, 81]. However, the molecular details of how these proteins undergo oscillatory dynamics are not clear. At each cell end, Cdc42 activation occurs via positive feedback and a time-delayed negative feedback resulting in the oscillatory pattern [11, 17, 46, 63, 67, 68, 82, 83]. The molecular details of how each cell-end overcomes this negative-feedback to allow reactivation of Cdc42 at that end is not known. Here we find that endocytosis enables periodic reactivation of Cdc42 at the cell ends by removing the negative-feedback and allowing the Cdc42 GEFs to return to that end.

Arp2/3 dependent endocytosis is known to be required for growth in *S. pombe* and other fungal species as well as mammalian cells with high membrane tension [75, 84, 85]. Previous works show that Arp2/3 dependent endocytosis is required to overcome the high internal turgor pressure for proper polarization and growth in yeasts [84, 86]. Here, we expound upon the importance of Arp2/3 dependent endocytosis for proper spatiotemporal regulation of polarity factors. We show that adequate removal of inhibitors is critical for proper polarized growth. Indeed, our lab has previously reported the importance of Rga4 removal for proper cell cycle progression and resumption of growth [87]. While previous works have described how exocytosis dilutes polarity protein concentration and dampens the concentration of active Cdc42 at that site, our research finds that endocytosis promotes Cdc42 activity through the removal of inhibition [52, 88].

The time-delayed negative feedback in Cdc42 regulation is mediated by its effector kinase Pak1 [11, 17, 26, 29, 30, 36]. Here we show that impeding endocytosis either by inhibition of the Arp2/3 complex or by deletion of *myo1* results in Pak1 accumulation at the cell ends. This indicates that endocytosis promotes Pak1 removal from the cell ends. Pak1 removal enables Scd1 localization to the cell ends and allows for proper Cdc42 activation dynamics between the two cell ends. Thus, endocytosis is required for maintaining anticorrelated Cdc42 activation dynamics between the cell ends thus promoting bipolar growth. The previous models have demonstrated the need for negative regulation on active Cdc42 for its oscillations [11, 13, 26, 68]. However, these models did not explore the impact of endocytosis on active Cdc42 dynamics and cell polarity. In this study, we developed models to illustrate how the accumulation of active Cdc42 hinders the buildup of Scd1 through the excess accumulation of Pak1 on the cellular membrane. Additionally, we found that endocytosis allows cellular bipolarity by facilitating the removal of Pak1.

In fission yeast, anticorrelated Cdc42 activation allows bipolar growth [11]. The cell end that exists from the previous generation is the old end while the end formed as a result of cell division is the new end. In fission yeast, the old end is the dominant end and will initiate Cdc42 activation first when daughter cells resume growth after cell division [9]. Bipolarity is established when the new end is able to overcome the old end dominance and robustly activate Cdc42 [9]. Anticorrelated Cdc42 activation between the two ends indicate that these ends compete for active Cdc42 or its regulators. Due to this competition, bipolar growth requires each end to undergo Cdc42 inactivation via negative feedback such that the opposite end can then activate Cdc42. Thus, each end must cycle between positive and negative feedback to allow bipolar Cdc42 activation.

Here we find that endocytosis has a role in maintaining the balance between positive feedback and time-delayed negative feedback to allow bipolar growth. While active Cdc42 is required for polarized growth, it is possible to have too much of a good thing. Absence of negative feedback, due to the loss of Pak1 activity, leads to an abundance of positive feedback at the dominant end. Thereby, Cdc42 activation at the dominant end is too strong for the second end to compete against, resulting in aberrant rounded morphology and monopolar growth as observed in *pak1-ts* mutants [11, 89]. We find that inhibition of the Arp2/3 complex due to CK-666 treatment or *myo1Δ* leads to enhanced negative feedback due to stabilization of Pak1 at the cell ends. This enhanced negative feedback results in monopolar Cdc42 activity and growth is inhibited at the site where Pak1 stabilizes. Positive feedback leads to Cdc42 activation which in turn allows Pak1 activation and time-delayed negative feedback. Pak1 has been shown to promote endocytosis (Fig.S3C) [70]. Our findings show that removal of negative feedback by Pak1-dependent endocytosis enables the return of positive feedback to the same cell end.

*pak1-ts* cells are characteristically monopolar. However, we observe that *pak1-ts* cells can localize Scd1 in a bipolar manner in a fraction of the cells, upon CK-666 treatment. We hypothesize that this may be explained by a secondary form of negative regulation mediated by Arp2/3 complex that is still unknown. As per Model 3, bipolar Scd1 in the absence of Pak1 occurs when Scd1 at the end is stabilized. It is possible that endocytosis partially contributes to Scd1 removal by an unknown mechanism and this is observable only in the absence of Pak1 kinase. Further research will investigate the mechanism of this second negative regulation. One possibility could be that endocytosis leads to changes in the levels of proteins at the cortex thus altering Cdc42 activation. Alternately, endocytosis may alter the lipid composition of the plasma membrane thereby affecting Cdc42 activation.

How are the proteins that participate in these multiple feedback loops spatiotemporally organized? While the functions and localizations of Cdc42 and its regulators have been extensively studied, the spatiotemporal localization (phase shifts) among these proteins had not been shown *in vivo*. Previous work has elucidated the phase shifts among polarity factors, ion signaling, and growth in pollen tubes [83, 90, 91]. In vitro experiments using *Xenopus* frog egg extracts and protein reconstituted systems have also shown phase shifts between waves of Rho GTPase activity and actin polymerization [46, 80]. Due to the advancements in live cell imaging and fluorophores, we have shown these phase shifts, *in vivo*, using the bipolar yeast, *S. pombe*. Our model predictions as well as experimental data show that the accumulation of active Cdc42 is in phase with the recruitment of its activator Scd1 but, by contrast, the accumulation of Pak1 shows a phase delay relative to the positive regulators. The delay between the Cdc42 and Pak1 accumulation further confirms the role of Pak1 as a time-delayed inhibitor of Cdc42 activity. The phase shift between Pak1 and the positive regulators likely results from its multiple roles and step wise regulation. First, Pak1 is only sufficiently recruited to the membrane once Cdc42 activation reaches a critical threshold ^15,16^. Next, it is likely that, Pak1 phosphorylates Scd1 to hinder Scd1-mediated Cd42 activation and dampen positive-feedback similar to budding yeast [35, 36]. Further, Pak1 plays multiple roles at the membrane in addition to Scd1 phosphorylation such as activating Myo1 which promotes Pak1 removal [32–34, 70]. Thus, the inhibitory role of increasing Pak1 activity causes a gradual decrease in its own recruitment until, eventually, Pak1 removal rate exceeds recruitment rate. This delay is further evidence of the complexity of self-organized processes and the multiple mechanisms in cell polarity.

Self-organization is seen throughout biology as a means to precisely regulate cellular processes [92]. Our work furthers the understanding of Cdc42’s extensive self-organizing capabilities [18, 93, 94]. Cdc42 activation recruits Scd2 and initiates positive feedback loops to activate more Cdc42 while simultaneously promoting Pak1 activation that triggers the negative feedback [11]. Cdc42 dynamics, its ability to accumulate at the membrane, and its potential to encounter its GAPs, are dependent on Cdc42 activation states [4, 13, 94, 95]. Likewise, Cdc42 activation also promotes actin polymerization for endocytosis and exocytosis [18, 40, 48, 50, 95–99]. We find that actin-mediated endocytosis is required for the removal of Pak1 to allow for Cdc42 activation and, as a result, both Pak1 recruitment and removal are facilitated by Cdc42. By the same token, continual Cdc42 activation and actin-mediated processes depend on endocytic removal of Pak1. Our findings show that the Cdc42 regulatory complex along with actin-mediated endocytosis form a self-organizing unit in the regulation of cell polarity.

In nature, robustness is important but cannot come at the expense of adaptability or vice versa. Complex signaling pathways with higher order molecular feedbacks have been observed in several processes and systems [92, 100–103]. These higher order pathways allow for adaptability while maintaining robustness. The RHO family GTPases undergo higher-order pathways via multiple feedback loops and this allows for their precise spatiotemporal activation necessary for cell polarization [6, 18, 104–107]. Our findings on Cdc42 regulation define higher order signaling pathways with multiple feedback loops. This regulation ensures polarized growth at each cell end and allows for bipolarity. While the role for Cdc42 regulating membrane trafficking is well documented, here we show how membrane trafficking also contributes to Cdc42 regulation and maintenance of bipolar growth.

## AUTHOR CONTRIBUTIONS

Conceptualization: M.A.H., Z.L., T.H., M.D.; Methodology: M.A.H., Z.L., T.H., M.D.; Validation: M.A.H., Z.L., B.F.C.,T.H.; Formal analysis: M.A.H., Z.L., B.F.C., L.C.; Investigation: M.A.H., Z.L., B.F.C.; Software: Z.L.; Resources: T.H., M.D.; Data curation: M.A.H.; Writing - original draft M.A.H.; Writing - review & editing: M.A.H., Z.L., B.F.C., T.H., M.D.; Visualization: M.A.H., Z.L., B.F.C.,T.H.; Supervision: T.H., M.D.; Project administration: M.D.; Funding acquisition: T.H., M.D.

## ACKNOWLEDGEMENTS

We thank James Moseley and Vladimir Sirotkin for providing strains and Bret Judson at Boston College Imaging core for imaging support.

---

**Table 1.**
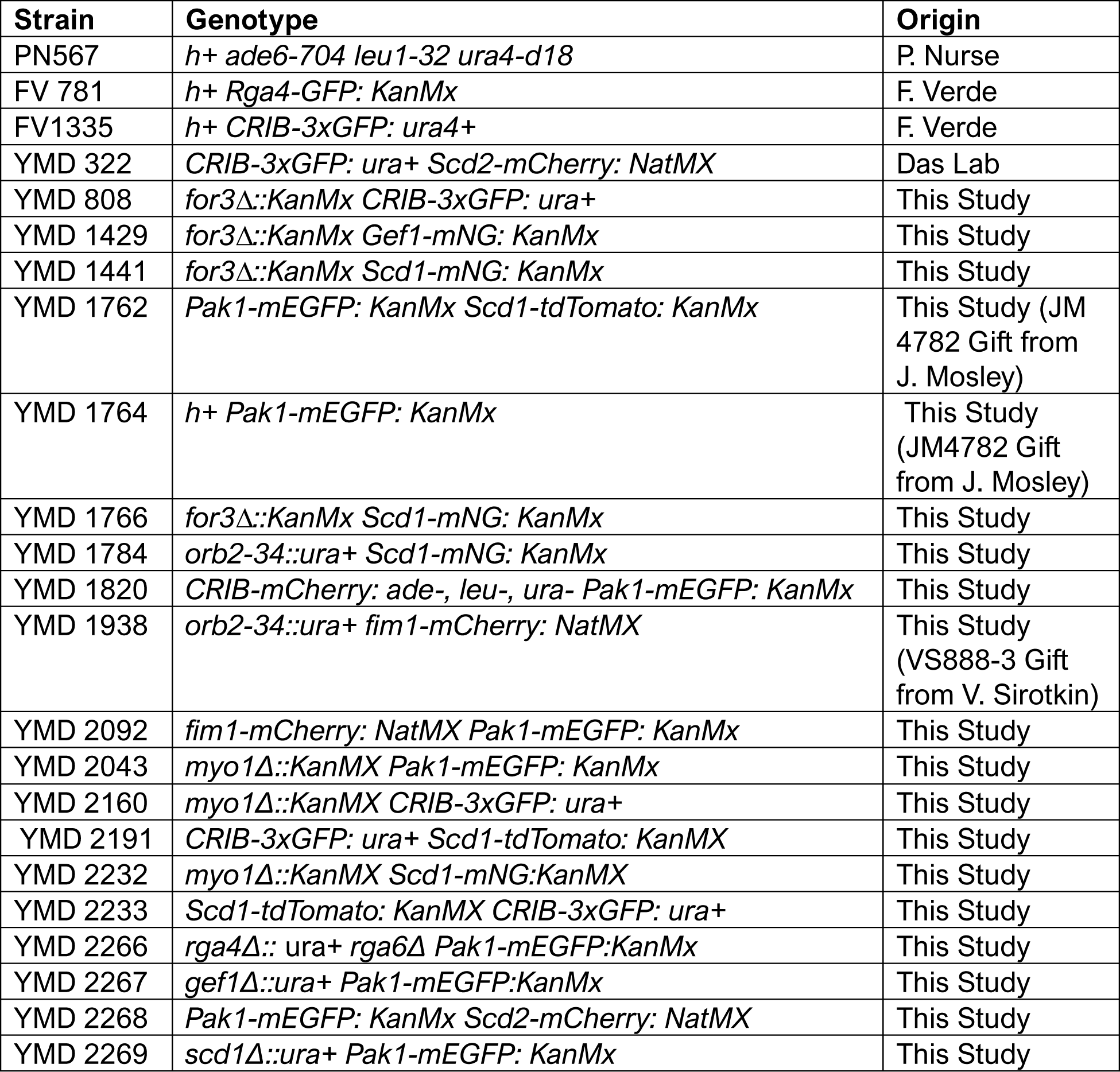
Strain list.

## STAR methods text

### Strains and Cell Culture

The S. pombe strains used in this study are listed in Table S1. All strains are isogenic to the original strain PN567. Cells were cultured in yeast extract (YE) medium and grown exponentially at 25°C, unless specified otherwise. Standard techniques were used for genetic manipulation and analysis [108]. Cells were grown exponentially for at least three rounds of eight generations each before imaging.

### Microscopy

Imaging was performed at room temperature (23–25°C). We used an Olympus IX83 microscope equipped with a VTHawk two-dimensional array laser scanning confocal microscopy system (Visitech International, Sunderland, UK), a Hamamatsu electron-multiplying charge-coupled device digital camera (Hamamatsu EM-CCD Digital Camera ImageM Model: C9100-13 Serial No: 741262, Hamamatsu, Hamamatsu City, Japan) and a 100×/1.49 NA UAPO lens (Olympus, Tokyo, Japan). Images were acquired using MetaMorph (Molecular Devices, Sunnyvale, CA).

We also used a spinning disk confocal microscope system with a Nikon Eclipse inverted microscope with a 100×/1.49 NA lens, a CSU-22 spinning disk system (Yokogawa Electric Corporation) and a Photometrics EM-CCD camera (Photometrics Technology Evolve with excelon Serial No: A13B107000). Images were acquired using MetaMorph (Molecular Devices, Sunnyvale, CA).

Additionally, we used a Nikon Ti2 Eclipse wide field microscope with a 100x/1.49 NA objective, an ORCA-FusionBT digital camera (Hamamatsu Model: C15440-20UP Serial No: 500428, Hamamatsu, Hamamatsu City, Japan). Images were acquired using Nikon NIS Elements (Nikon, Melville, NY). Fluorophores were excited using an AURA Light Engine system (Lumencor, Beaverton, OR).

Microscopy was also performed with a 3i spinning disk confocal using a Zeiss AxioObserver microscope with integrated Yokogawa spinning disk (Yokogawa CSU-X1 A1 spinning disk scanner) and a 100x/1.49 NA objective. Images were acquired with a Teledyne Photometrics Prime 95b back-illuminated sCMOS camera (Serial No: A20D203014, Tucson, AZ). Images were acquired using SlideBook (3i Intelligent Imaging innovations, Denver, CO).

### Acquiring and quantifying fluorescence Intensity

Fluorescence intensity was measured using ImageJ software. All images were sum projected and mean intensities were reported. Freehand ROIs were used to measure signal at the cell ends. For analysis of two ends of the same cell, cytoplasm with little or no signal was used for background subtraction. For all other experiments, a region outside of the cell was used for background subtraction. The mean intensity was measured and recorded after the background subtractions.

To quantify the anticorrelation between two ends, cells in each condition were imaged every minute for 60 minutes. The data from each cell was then analyzed to find the correlation coefficient between fluorescent signals of both ends within the cell.

To quantify signal stability at the cell ends, cells were imaged every 30 seconds for 20 minutes. The stability of the proteins was then quantified by measuring the difference in intensity between each subsequent timepoint for the duration of the timelapse imaging. Populations that had larger average variations of fluorescent intensity between each timepoint were more dynamic and less stable than populations that had smaller variations between each timepoint.

To observe and quantify the in vivo phase-shifts, cells were imaged every 6 seconds for 9 minutes. Two fluorescent proteins were imaged simultaneously using the 488 nm and 561 nm wavelengths. LED power was set to 3% for each wavelength. Standard fluorescence intensity method was used to measure each timepoint. The phase-lag between each protein at each cell end was then quantified using cross-correlation analysis wherein the correlation coefficient is calculated for both signals with and without shifting the quantified signal. The second fluorescent signal is shifted backwards by 20 timepoints while the first signal remains the same and the correlation coefficient is calculated for each shift. The correlation coefficient is also calculated as the signal is shifted forward by 20 timepoints and calculated for each shift. The number of shifts required to achieve the best correlation coefficient indicates how far one signal lags the other. The shift required to produce the best correlation between the two fluorescent signals was verified as showing true correlation or not before being included in the final comparative analysis for each set of protein signals.

To observe the loss of Pak1 from the cell membrane in relation to the internalization of Fim1, cells were imaged each second for 3 minutes. Square ROIs were used to isolate regions of the cell end for analysis. These ROIs were used to make kymographs of both Fim1-mCherry signal and Pak1m-EGFP signal. These kymographs were quantified by plotting the intensity profile of both proteins at the cell membrane. These intensity profiles were then analyzed by recording when Pak1 was lost in comparison to each successful endocytic event. Between 6-10 endocytic events were captured in each kymograph. These events were then averaged for each kymograph and graphed.

### Actin Cytoskeleton disruptions

Cells were treated with 100 µM CK-666 (Sigma-Aldrich, SML006-5MG) in DMSO (Sigma-Aldrich, D8418-250ML) to block the Arp2/3 complex and branched actin assembly. To block the polymerization of all F-actin structures, cells were treated with 10 µM Latrunculin A (LatA; EMD Millipore) dissolved in DMSO for 30 minutes prior to imaging. Control cells were treated with 0.1% DMSO in YE media.

### Statistical Tests

Significance was determined using GraphPad Prism. When comparing two conditions, a Student’s t-test was used (two-tailed, unequal variance). One way ANOVA, followed by a Tukey’s multiple comparison test, was used to determine significance for experiments with three or more conditions.

### Mathematical models for regulation networks

The details of the regulatory network are provided in the supplementary materials. This paper introduces three networks, all described using a reaction-diffusion model based on the Xu-Jilkine model [68]. Model 1 and Model 2 each include one negative regulatory pathway of active Cdc42. In Model 1, there is a direct inhibition from active Cdc42 to Scd1. In Model 2, the negative pathway involves Pak1. Model 3, on the other hand, includes two negative regulatory pathways of active Cdc42, with one involving Pak1. However, the details of the second pathway remain unclear. For the numerical analysis, MATLAB was used.

### Computational Phase analysis

Z-scores were used to normalize the data. Prior to analysis, the normalized data was smoothened, and the Hilbert Transform of the data was applied using MATLAB.

## REFERENCES

1. Drubin, D.G., and Nelson, W.J. (1996). Origins of Cell Polarity. Cell 84, 335–344.

2. Glotzer, M., and Hyman, A.A. (1995). Cell Polarity: The importance of being polar. Current Biology 5, 1102–1105.

3. Nobes, C.D., and Hall, A. (1999). Rho GTPases control polarity, protrusion, and adhesion during cell movement. J Cell Biol 144, 1235–1244.

4. Etienne-Manneville, S. (2004). Cdc42--the centre of polarity. J Cell Sci 117, 1291–1300.

5. Manser, E., Leung, T., Salihuddin, H., Zhao, Z.S., and Lim, L. (1994). A brain serine/threonine protein kinase activated by Cdc42 and Rac1. Nature 367, 40–46.

6. Martin-Vilchez, S., Whitmore, L., Asmussen, H., Zareno, J., Horwitz, R., and Newell-Litwa, K. (2017). RhoGTPase Regulators Orchestrate Distinct Stages of Synaptic Development. PloS one 12, e0170464.

7. Machacek, M., Hodgson, L., Welch, C., Elliott, H., Pertz, O., Nalbant, P., Abell, A., Johnson, G.L., Hahn, K.M., and Danuser, G. (2009). Coordination of Rho GTPase activities during cell protrusion. Nature 461, 99–103.

8. de Beco, S., Vaidziulyte, K., Manzi, J., Dalier, F., di Federico, F., Cornilleau, G., Dahan, M., and Coppey, M. (2018). Optogenetic dissection of Rac1 and Cdc42 gradient shaping. Nat Commun 9, 4816.

9. Mitchison, J.M., and Nurse, P. (1985). Growth in cell length in the fission yeast Schizosaccharomyces pombe. J Cell Sci 75, 357–376.

10. Johnson, D.I. (1999). Cdc42: An essential Rho-type GTPase controlling eukaryotic cell polarity. Microbiol Mol Biol Rev 63, 54–105.

11. Das, M., Drake, T., Wiley, D.J., Buchwald, P., Vavylonis, D., and Verde, F. (2012). Oscillatory dynamics of Cdc42 GTPase in the control of polarized growth. Science 337, 239–243.

12. Butty, A.C. (2002). A positive feedback loop stabilizes the guanine-nucleotide exchange factor Cdc24 at sites of polarization. The EMBO Journal 21, 1565–1576.

13. Das, M., and Verde, F. (2013). Role of Cdc42 dynamics in the control of fission yeast cell polarization. Biochemical Society transactions 41, 1745–1749.

14. Turing, A.M. (1990). The chemical basis of morphogenesis. 1953. Bull Math Biol 52, 153–197; discussion 119-152.

15. Wu, C.F., Chiou, J.G., Minakova, M., Woods, B., Tsygankov, D., Zyla, T.R., Savage, N.S., Elston, T.C., and Lew, D.J. (2015). Role of competition between polarity sites in establishing a unique front. Elife 4.

16. Chiou, J.G., Ramirez, S.A., Elston, T.C., Witelski, T.P., Schaeffer, D.G., and Lew, D.J. (2018). Principles that govern competition or co-existence in Rho-GTPase driven polarization. PLoS Comput Biol 14, e1006095.

17. Wu, C.F., and Lew, D.J. (2013). Beyond symmetry-breaking: competition and negative feedback in GTPase regulation. Trends in cell biology 23, 476–483.

18. Lamas, I., Merlini, L., Vjestica, A., Vincenzetti, V., and Martin, S.G. (2020). Optogenetics reveals Cdc42 local activation by scaffold-mediated positive feedback and Ras GTPase. PLoS biology 18, e3000600.

19. Hercyk, B.S., Rich-Robinson, J., Mitoubsi, A.S., Harrell, M.A., and Das, M.E. (2019). A novel interplay between GEFs orchestrates Cdc42 activity during cell polarity and cytokinesis. Journal of Cell Science 132, jcs236018.

20. Chang, E.C., Barr, M., Wang, Y., Jung, V., Xu, H.P., and Wigler, M.H. (1994). Cooperative interaction of S. pombe proteins required for mating and morphogenesis. Cell 79, 131–141.

21. Endo, M., Shirouzu, M., and Yokoyama, S. (2003). The Cdc42 binding and scaffolding activities of the fission yeast adaptor protein Scd2. The Journal of biological chemistry 278, 843–852.

22. Revilla-Guarinos, M.T., Martín-García, R., Villar-Tajadura, M.A., Estravís, M., Coll, P.M., and Pérez, P. (2016). Rga6 is a Fission Yeast Rho GAP Involved in Cdc42 Regulation of Polarized Growth. Mol Biol Cell 27, 1524–1535.

23. Das, M., Wiley, D.J., Medina, S., Vincent, H.A., Larrea, M., Oriolo, A., and Verde, F. (2007). Regulation of cell diameter, For3p localization, and cell symmetry by fission yeast Rho-GAP Rga4p. Mol Biol Cell 18, 2090–2101.

24. Gallo Castro, D., and Martin, S.G. (2018). Differential GAP requirement for Cdc42-GTP polarization during proliferation and sexual reproduction. The Journal of cell biology 217, 4215–4229.

25. Tatebe, H., Nakano, K., Maximo, R., and Shiozaki, K. (2008). Pom1 DYRK regulates localization of the Rga4 GAP to ensure bipolar activation of Cdc42 in fission yeast. Curr Biol 18, 322–330.

26. Howell, A.S., Jin, M., Wu, C.F., Zyla, T.R., Elston, T.C., and Lew, D.J. (2012). Negative feedback enhances robustness in the yeast polarity establishment circuit. Cell 149, 322–333.

27. McCusker, D., and Kellogg, D.R. (2012). Plasma membrane growth during the cell cycle: unsolved mysteries and recent progress. Current opinion in cell biology 24, 845–851.

28. Marcus, S., Polverino, A., Chang, E., Robbins, D., Cobb, M.H., and Wigler, M.H. (1995). Shk1, a homolog of the Saccharomyces cerevisiae Ste20 and mammalian p65PAK protein kinases, is a component of a Ras/Cdc42 signaling module in the fission yeast Schizosaccharomyces pombe. Proceedings of the National Academy of Sciences 92, 6180–6184.

29. Tu, H., and Wigler, M. (1999). Genetic evidence for Pak1 autoinhibition and its release by Cdc42. Molecular and cellular biology 19, 602–611.

30. Ottilie, S., Miller, P.J., Johnson, D.I., Creasy, C.L., Sells, M.A., Bagrodia, S., Forsburg, S.L., and Chernoff, J. (1995). Fission yeast pak1+ encodes a protein kinase that interacts with Cdc42p and is involved in the control of cell polarity and mating. The EMBO Journal 14, 5908–5919.

31. Molli, P.R., Li, D.Q., Murray, B.W., Rayala, S.K., and Kumar, R. (2009). PAK signaling in oncogenesis. Oncogene 28, 2545–2555.

32. Onwubiko, U.N., Kalathil, D., Koory, E., Pokharel, S., Roberts, H., Mitoubsi, A., and Das, M. (2023). Cdc42 prevents precocious Rho1 activation during cytokinesis in a Pak1-dependent manner. J Cell Sci 136.

33. Magliozzi, J.O., Sears, J., Cressey, L., Brady, M., Opalko, H.E., Kettenbach, A.N., and Moseley, J.B. (2020). Fission yeast Pak1 phosphorylates anillin-like Mid1 for spatial control of cytokinesis. J Cell Biol 219.

34. Magliozzi, J.O., and Moseley, J.B. (2021). Pak1 kinase controls cell shape through ribonucleoprotein granules. Elife 10.

35. Kuo, C.C., Savage, N.S., Chen, H., Wu, C.F., Zyla, T.R., and Lew, D.J. (2014). Inhibitory GEF phosphorylation provides negative feedback in the yeast polarity circuit. Current biology : CB 24, 753–759.

36. Rapali, P., Mitteau, R., Braun, C., Massoni-Laporte, A., Unlu, C., Bataille, L., Arramon, F.S., Gygi, S.P., and McCusker, D. (2017). Scaffold-mediated gating of Cdc42 signalling flux. Elife 6.

37. Das, M., Wiley, D.J., Chen, X., Shah, K., and Verde, F. (2009). The Conserved NDR Kinase Orb6 Controls Polarized Cell Growth by Spatial Regulation of the Small GTPase Cdc42. Current Biology 19, 1314–1319.

38. Kolluri, R., Tolias, K.F., Carpenter, C.L., Rosen, F.S., and Kirchhausen, T. (1996). Direct interaction of the Wiskott-Aldrich syndrome protein with the GTPase Cdc42. Proc Natl Acad Sci U S A 93, 5615–5618.

39. Lechler, T., Shevchenko, A., and Li, R. (2000). Direct involvement of yeast type I myosins in Cdc42-dependent actin polymerization. The Journal of cell biology 148, 363–373.

40. Kovar, D.R., Sirotkin, V., and Lord, M. (2011). Three’s company: the fission yeast actin cytoskeleton. Trends Cell Biol 21, 177–187.

41. Onwubiko, U.N., Rich-Robinson, J., Mustaf, R.A., and Das, M.E. (2021). Cdc42 promotes Bgs1 recruitment for septum synthesis and glucanase localization for cell separation during cytokinesis in fission yeast. Small GTPases 12, 257–264.

42. Onwubiko, U.N., Mlynarczyk, P.J., Wei, B., Habiyaremye, J., Clack, A., Abel, S.M., and Das, M.E. (2019). A Cdc42 GEF, Gef1, through endocytosis organizes F-BAR Cdc15 along the actomyosin ring and promotes concentric furrowing. Journal of cell science 132.

43. Thomas, Jenna, Barone, E., Suarez, C., Sirotkin, V., and David (2014). Homeostatic Actin Cytoskeleton Networks Are Regulated by Assembly Factor Competition for Monomers. Current Biology 24, 579–585.

44. Bendezú, F.O., and Martin, S.G. (2011). Actin cables and the exocyst form two independent morphogenesis pathways in the fission yeast. Molecular Biology of the Cell 22, 44–53.

45. Martin, S.G., Rincon, S.A., Basu, R., Perez, P., and Chang, F. (2007). Regulation of the formin for3p by cdc42p and bud6p. Molecular biology of the cell 18, 4155–4167.

46. Landino, J., Leda, M., Michaud, A., Swider, Z.T., Prom, M., Field, C.M., Bement, W.M., Vecchiarelli, A.G., Goryachev, A.B., and Miller, A.L. (2021). Rho and F-actin self-organize within an artificial cell cortex. Curr Biol 31, 5613–5621 e5615.

47. Lee, W.-L., Bezanilla, M., and Pollard, T.D. (2000). Fission Yeast Myosin-I, Myo1p, Stimulates Actin Assembly by Arp2/3 Complex and Shares Functions with Wasp. Journal of Cell Biology 151, 789–800.

48. Gachet, Y., and Hyams, J.S. (2005). Endocytosis in fission yeast is spatially associated with the actin cytoskeleton during polarised cell growth and cytokinesis. Journal of cell science 118, 4231–4242.

49. Sirotkin, V., Beltzner, C.C., Marchand, J.B., and Pollard, T.D. (2005). Interactions of WASp, myosin-I, and verprolin with Arp2/3 complex during actin patch assembly in fission yeast. The Journal of cell biology 170, 637–648.

50. Campbell, B.F., Hercyk, B.S., Williams, A.R., San Miguel, E., Young, H.G., and Das, M.E. (2022). Cdc42 GTPase activating proteins Rga4 and Rga6 coordinate septum synthesis and membrane trafficking at the division plane during cytokinesis. Traffic 23, 478–495.

51. Hercyk, B.S., and Das, M.E. (2019). F-BAR Cdc15 Promotes Cdc42 Activation During Cytokinesis and Cell Polarization in Schizosaccharomyces pombe. Genetics 213, 1341–1356.

52. Watson, L.J., Rossi, G., and Brennwald, P. (2014). Quantitative Analysis of Membrane Trafficking in Regulation of Cdc42 Polarity. Traffic 15, 1330–1343.

53. Pelham, R.J., and Chang, F. (2001). Role of actin polymerization and actin cables in actin-patch movement in Schizosaccharomyces pombe. Nature Cell Biology 3, 235–244.

54. Salat-Canela, C., Carmona, M., Martín-García, R., Pérez, P., Ayté, J., and Hidalgo, E. (2021). Stress-dependent inhibition of polarized cell growth through unbalancing the GEF/GAP regulation of Cdc42. Cell Rep 37, 109951.

55. Mutavchiev, D.R., Leda, M., and Sawin, K.E. (2016). Remodeling of the Fission Yeast Cdc42 Cell-Polarity Module via the Sty1 p38 Stress-Activated Protein Kinase Pathway. Curr Biol 26, 2921–2928.

56. Fujiwara, I., Zweifel, M.E., Courtemanche, N., and Pollard, T.D. (2018). Latrunculin A Accelerates Actin Filament Depolymerization in Addition to Sequestering Actin Monomers. Curr Biol 28, 3183–3192.e3182.

57. Morton, W.M., Ayscough, K.R., and McLaughlin, P.J. (2000). Latrunculin alters the actin-monomer subunit interface to prevent polymerization. Nat Cell Biol 2, 376–378.

58. Beltzner, C.C., and Pollard, T.D. (2008). Pathway of actin filament branch formation by Arp2/3 complex. J Biol Chem 283, 7135–7144.

59. Feierbach, B., and Chang, F. (2001). Roles of the fission yeast formin for3p in cell polarity, actin cable formation and symmetric cell division. Curr Biol 11, 1656–1665.

60. Martin, S.G., and Chang, F. (2006). Dynamics of the formin for3p in actin cable assembly. Curr Biol 16, 1161–1170.

61. Pelham, R.J., and Chang, F. (2002). Actin dynamics in the contractile ring during cytokinesis in fission yeast. Nature 419, 82–86.

62. Nolen, B.J., Tomasevic, N., Russell, A., Pierce, D.W., Jia, Z., McCormick, C.D., Hartman, J., Sakowicz, R., and Pollard, T.D. (2009). Characterization of two classes of small molecule inhibitors of Arp2/3 complex. Nature 460, 1031–1034.

63. Das, M., Nunez, I., Rodriguez, M., Wiley, D.J., Rodriguez, J., Sarkeshik, A., Yates, J.R., 3rd, Buchwald, P., and Verde, F. (2015). Phosphorylation-dependent inhibition of Cdc42 GEF Gef1 by 14-3-3 protein Rad24 spatially regulates Cdc42 GTPase activity and oscillatory dynamics during cell morphogenesis. Mol Biol Cell 26, 3520–3534.

64. Stamnes, M. (2002). Regulating the actin cytoskeleton during vesicular transport. Curr Opin Cell Biol 14, 428–433.

65. Gundelfinger, E.D., Kessels, M.M., and Qualmann, B. (2003). Temporal and spatial coordination of exocytosis and endocytosis. Nature reviews. Molecular cell biology 4, 127–139.

66. Wu, L.G., Hamid, E., Shin, W., and Chiang, H.C. (2014). Exocytosis and endocytosis: modes, functions, and coupling mechanisms. Annu Rev Physiol 76, 301–331.

67. Novák, B., and Tyson, J.J. (2008). Design principles of biochemical oscillators. Nature Reviews Molecular Cell Biology 9, 981–991.

68. Xu, B., and Jilkine, A. (2018). Modeling the Dynamics of Cdc42 Oscillation in Fission Yeast. Biophysical journal 114, 2025.

69. Goodwin, B.C. (1965). Oscillatory behavior in enzymatic control processes. Adv Enzyme Regul 3, 425–438.

70. Attanapola, S.L., Alexander, C.J., and Mulvihill, D.P. (2009). Ste20-kinase-dependent TEDS-site phosphorylation modulates the dynamic localisation and endocytic function of the fission yeast class I myosin, Myo1. Journal of Cell Science 122, 3856–3861.

71. Grassart, A., Dujeancourt, A., Lazarow, P.B., Dautry-Varsat, A., and Sauvonnet, N. (2008). Clathrin-independent endocytosis used by the IL-2 receptor is regulated by Rac1, Pak1 and Pak2. EMBO Rep 9, 356–362.

72. Nakano, K., Satoh, K., Morimatsu, A., Ohnuma, M., and Mabuchi, I. (2001). Interactions among a fimbrin, a capping protein, and an actin-depolymerizing factor in organization of the fission yeast actin cytoskeleton. Mol Biol Cell 12, 3515–3526.

73. Macquarrie, C.D., Mangione, M.C., Carroll, R., James, M., Gould, K.L., and Sirotkin, V. (2019). Adaptor protein Bbc1 regulates localization of Wsp1 and Vrp1 during endocytic actin patch assembly. Journal of Cell Science 132, jcs233502.

74. Arasada, R., and Pollard, Thomas D. (2011). Distinct Roles for F-BAR Proteins Cdc15p and Bzz1p in Actin Polymerization at Sites of Endocytosis in Fission Yeast. Current Biology 21, 1450–1459.

75. Basu, R., Munteanu, E.L., and Chang, F. (2014). Role of turgor pressure in endocytosis in fission yeast. Molecular biology of the cell 25, 679–687.

76. Cao, Y., Lopatkin, A., and You, L. (2016). Elements of biological oscillations in time and space. Nat Struct Mol Biol 23, 1030–1034.

77. Wheatley, E., and Rittinger, K. (2005). Interactions between Cdc42 and the scaffold protein Scd2: requirement of SH3 domains for GTPase binding. The Biochemical journal 388, 177–184.

78. Lamas, I., Weber, N., and Martin, S.G. (2020). Activation of Cdc42 GTPase upon CRY2-Induced Cortical Recruitment Is Antagonized by GAPs in Fission Yeast. Cells 9.

79. Sabherwal, N., Rowntree, A., Marinopoulou, E., Pettini, T., Hourihane, S., Thomas, R., Soto, X., Kursawe, J., and Papalopulu, N. (2021). Differential phase register of Hes1 oscillations with mitoses underlies cell-cycle heterogeneity in ER(+) breast cancer cells. Proc Natl Acad Sci U S A 118.

80. Bement, W.M., Leda, M., Moe, A.M., Kita, A.M., Larson, M.E., Golding, A.E., Pfeuti, C., Su, K.C., Miller, A.L., Goryachev, A.B., et al. (2015). Activator-inhibitor coupling between Rho signalling and actin assembly makes the cell cortex an excitable medium. Nat Cell Biol 17, 1471–1483.

81. Coll, P.M., Trillo, Y., Ametzazurra, A., and Perez, P. (2003). Gef1p, a new guanine nucleotide exchange factor for Cdc42p, regulates polarity in Schizosaccharomyces pombe. Mol Biol Cell 14, 313–323.

82. Tsai, T.Y.-C., Choi, Y.S., Ma, W., Pomerening, J.R., Tang, C., and Ferrell, J.E. (2008). Robust, Tunable Biological Oscillations from Interlinked Positive and Negative Feedback Loops. Science 321, 126–129.

83. Hwang, J.-U., Gu, Y., Lee, Y.-J., and Yang, Z. (2005). Oscillatory ROP GTPase Activation Leads the Oscillatory Polarized Growth of Pollen Tubes. Molecular Biology of the Cell 16, 5385–5399.

84. Aghamohammadzadeh, S., and Ayscough, K.R. (2009). Differential requirements for actin during yeast and mammalian endocytosis. Nat Cell Biol 11, 1039–1042.

85. Epp, E., Walther, A., Lépine, G., Leon, Z., Mullick, A., Raymond, M., Wendland, J., and Whiteway, M. (2010). Forward genetics in Candida albicans that reveals the Arp2/3 complex is required for hyphal formation, but not endocytosis. Mol Microbiol 75, 1182–1198.

86. Basu, R., Munteanu, E.L., and Chang, F. (2014). Role of turgor pressure in endocytosis in fission yeast. Molecular Biology of the Cell 25, 679–687.

87. Rich-Robinson, J., Russell, A., Mancini, E., and Das, M. (2021). Cdc42 reactivation at growth sites is regulated by local cell-cycle-dependent loss of its GTPase-activating protein Rga4 in fission yeast. Journal of cell science 134.

88. Ghose, D., and Lew, D. (2020). Mechanistic insights into actin-driven polarity site movement in yeast. Mol Biol Cell 31, 1085–1102.

89. Sawin, K.E., Hajibagheri, M.A.N., and Nurse, P. (1999). Mis-specification of cortical identity in a fission yeast PAK mutant. Current Biology 9, 1335–1338.

90. Messerli, M., and Robinson, K.R. (1997). Tip localized Ca2+ pulses are coincident with peak pulsatile growth rates in pollen tubes of Lilium longiflorum. J Cell Sci 110 ( *Pt 11*), 1269–1278.

91. Messerli, M.A., Danuser, G., and Robinson, K.R. (1999). Pulsatile influxes of H+, K+ and Ca2+ lag growth pulses of Lilium longiflorum pollen tubes. J Cell Sci 112 (*Pt 10*), 1497–1509.

92. Karsenti, E. (2008). Self-organization in cell biology: a brief history. Nature reviews. Molecular cell biology 9, 255–262.

93. Gerganova, V., Lamas, I., Rutkowski, D.M., Vještica, A., Castro, D.G., Vincenzetti, V., Vavylonis, D., and Martin, S.G. (2021). Cell patterning by secretion-induced plasma membrane flows. Sci Adv 7, eabg6718.

94. Rutkowski, D.M., Vincenzetti, V., Vavylonis, D., and Martin, S.G. (2023). Cdc42 mobility and membrane flows regulate fission yeast cell shape and survival. bioRxiv.

95. Estravís, M., Rincón, S.A., Portales, E., Pérez, P., and Santos, B. (2017). Cdc42 activation state affects its localization and protein levels in fission yeast. Microbiology 163, 1156–1166.

96. Sagot, I., Klee, S.K., and Pellman, D. (2002). Yeast formins regulate cell polarity by controlling the assembly of actin cables. Nat Cell Biol 4, 42–50.

97. Adamo, J.E., Moskow, J.J., Gladfelter, A.S., Viterbo, D., Lew, D.J., and Brennwald, P.J. (2001). Yeast Cdc42 functions at a late step in exocytosis, specifically during polarized growth of the emerging bud. J Cell Biol 155, 581–592.

98. Estravis, M., Rincon, S.A., Santos, B., and Perez, P. (2011). Cdc42 regulates multiple membrane traffic events in fission yeast. Traffic 12, 1744–1758.

99. Estravis, M., Rincon, S., and Perez, P. (2012). Cdc42 regulation of polarized traffic in fission yeast. Communicative & integrative biology 5, 370–373.

100. Glazenburg, M.M., and Laan, L. (2023). Complexity and self-organization in the evolution of cell polarization. J Cell Sci 136.

101. Guo, X., and Dong, J. (2022). Protein polarization: Spatiotemporal precisions in cell division and differentiation. Curr Opin Plant Biol 68, 102257.

102. Landge, A.N., Jordan, B.M., Diego, X., and Müller, P. (2020). Pattern formation mechanisms of self-organizing reaction-diffusion systems. Dev Biol 460, 2–11.

103. Stock, J., and Pauli, A. (2021). Self-organized cell migration across scales - from single cell movement to tissue formation. Development 148.

104. Nobes, C.D., and Hall, A. (1995). Rho, rac, and cdc42 GTPases regulate the assembly of multimolecular focal complexes associated with actin stress fibers, lamellipodia, and filopodia. Cell 81, 53–62.

105. Martin, K., Reimann, A., Fritz, R.D., Ryu, H., Jeon, N.L., and Pertz, O. (2016). Spatio-temporal co-ordination of RhoA, Rac1 and Cdc42 activation during prototypical edge protrusion and retraction dynamics. Sci Rep 6, 21901.

106. Fairn, G.D., Hermansson, M., Somerharju, P., and Grinstein, S. (2011). Phosphatidylserine is polarized and required for proper Cdc42 localization and for development of cell polarity. Nat Cell Biol 13, 1424–1430.

107. Hedrick, N.G., Harward, S.C., Hall, C.E., Murakoshi, H., McNamara, J.O., and Yasuda, R. (2016). Rho GTPase complementation underlies BDNF-dependent homo-and heterosynaptic plasticity. Nature 538, 104–108.

108. Moreno, S., Klar, A., and Nurse, P. (1991). Molecular genetic analysis of fission yeast Schizosaccharomyces pombe. Methods Enzymol 194, 795–823.

