## Supplemental material and figures for "The Arp2/3 complex promotes periodic removal of Pak1-mediated negative feedback to facilitate anticorrelated Cdc42 oscillations"

#### Mathematical Modeling

**Model 1:** The model is constructed using reaction-diffusion equations to describe the regulatory network. The network includes Cdc42-GTP, and Scd1 at the tips, as well as the diffusion of inactive molecules like Cdc42-GDP, and Scd1 in the cytoplasm.

The diffusion part in the cytoplasm is:

$$\frac{\partial C}{\partial t}(x,t) = D_C \frac{\partial^2 C}{\partial x^2}(x,t) \qquad (1)$$
$$\frac{\partial S}{\partial t}(x,t) = D_S \frac{\partial^2 S}{\partial x^2}(x,t). \qquad (2)$$

$D_C$ and $D_S$ are diffusion coefficients of Cdc42-GDP and Scd1, respectively.
The reaction part at tips is

$$\frac{dc_i}{dt}(t) = k^c(s_i(t))C(L_i,t) - \delta_c c_i(t), \quad i = 1,2 \qquad (4)$$
$$\frac{ds_i}{dt}(t) = k^s(c_i(t))S(L_i,t) - \delta_s s_i(t), \quad i = 1,2 \qquad (5)$$
$$k^c(s_i(t)) = \frac{k_c s_i(t)^{n_{sc}}}{s_i(t)^{n_{sc}} + K_{sc}{}^{n_{sc}}} \qquad (7)$$
$$k^s(c_i(t)) = \frac{k_s c_i(t)^{n_{cs}}}{(c_i(t)^{n_{cs}} + K_{cs}{}^{n_{cs}})(1 + \left(\frac{c_i(t)}{K_{as}}\right)^{n_{as}})}. \qquad (8)$$

$c_i$, $g_i$ and $p_i$ are active Cdc42, Scd1, and Pak1 at two tips. $\delta_C$ is the degradation rate constant for molecule Cdc42-GTP. $\delta_S$ is the degradation rate constant for molecule Scd1. One time unit is one minute.

**Model 2:** The model structure is similar to Model 1. The network includes Cdc42-GTP, protein X (Pak1), and Scd1 at the tips, as well as the diffusion of inactive molecules like Cdc42-GDP, protein X (Pak1), and Scd1 within the cytoplasm. Pak1 is described as protein X at first.

The diffusion part:

$$\frac{\partial C}{\partial t}(x,t) = D_C \frac{\partial^2 C}{\partial x^2}(x,t) \qquad (9)$$
$$\frac{\partial S}{\partial t}(x,t) = D_S \frac{\partial^2 S}{\partial x^2}(x,t) \qquad (10)$$
$$\frac{\partial P}{\partial t}(x,t) = D_P \frac{\partial^2 P}{\partial x^2}(x,t) \qquad (11)$$

$D_C$, $D_S$ and $D_P$ are diffusion coefficients of Cdc42-GDP, Scd1, and Pak1, respectively.

The equations at tips for the second model are:

$$\frac{dc_i}{dt}(t) = k^c(s_i(t))C(L_i,t) - \delta_c c_i(t), \quad i = 1,2 \qquad (12)$$
$$\frac{ds_i}{dt}(t) = k^s(c_i(t), p_i(t))S(L_i,t) - \delta_s s_i(t), \quad i = 1,2 \qquad (13)$$
$$\frac{dp_i}{dt}(t) = k^p\big(c_i(t)\big)P(L_i,t) - \delta_p p_i(t) \quad i = 1,2 \qquad (14)$$

$$k^c(s_i(t)) = \frac{k_c s_i(t)^{n_{sc}}}{s_i(t)^{n_{sc}} + K_{sc}{}^{n_{sc}}} \qquad (15)$$
$$k^s(c_i(t), p_i(t)) = \frac{k_s c_i(t)^{n_{cs}}}{(c_i(t)^{n_{cs}} + K_{cs}{}^{n_{cs}})(1 + (\frac{p_i(t)}{K_{ps}})^{n_{ps}})} \qquad (16)$$
$$k^p(c_i(t)) = \frac{k_p c_i(t)^{n_{cp}}}{c_i(t)^{n_{cp}} + K_{cp}{}^{n_{cp}}}. \qquad (17)$$

$c_i$, $g_i$ and $p_i$ are active Cdc42, Scd1, and Pak1 at two tips. $\delta_C$ and $\delta_S$ are the same in Model1, $\delta_p$ is the degradation rate constant for molecule Pak1.

**Model 3:** Incorporating another negative feedback loop into the Cdc42-GTP regulatory network, we seek to provide an explanation for bipolarity in the absence of Pak1. However, the specific details of this second feedback loop are unknown. As a result, we made the assumption that active Cdc42 can inhibit itself, either directly or indirectly. Eq 12 is changed to:

$$\frac{dc_i}{dt}(t) = k^c(s_i(t), c_i(t)\,)C(L_i, t) - \delta_c c_i(t), \quad i = 1{,}2 \qquad (12a)$$

and Eq 15 is changed to:

$$k^c(s_i(t), c_i(t)) = \frac{k_c s_i(t)^{n_{sc}}}{s_i(t)^{n_{sc}} + K_{sc}{}^{n_{sc}}}\ e^{\frac{-c_i(t)}{a1}} \quad . \qquad (15a)$$

Here $a1$ is a constant describing the strength of the second negative feedback loop.

**Comparison of oscillatory dynamics between Model 1 and Model 2:** In MATLAB, Sobol parameter spaces are defined for each model, with each model having 100,000 sample points in its parameter space. The objective is to detect oscillatory dynamics by counting peaks and measuring the differences in amplitude between neighboring peaks. The parameter sets that lead to oscillatory dynamics are counted and used to calculate the percentage of occurrences of oscillatory dynamics. (Range of parameters are in table S1)

**Model 2-1, Pak1 promotes endocytosis:** We change Eq 14 to

$$\delta_p = \delta_p' f(p)$$

$$f(p) = \frac{k_{pe} p_i(t)^{n_{pe}}}{p_i(t)^{n_{pe}} + K_{pe}{}^{n_{pe}}}.$$

Here $\delta_p'$ is a constant.

**Table S1. Parameter values of the wild-type model2, including parameters in model2-1.**

| Parameter | Description | Value | Unit | Range for oscillation search |
|---|---|---|---|---|
| $D_C$ | Diffusion of Cdc42-GDP in cytoplasm | 5 | a.u. length/s | (0,10) |
| $D_S$ | Diffusion of Scd1 in cytoplasm | 5 | a.u. length/s | (0,10) |
| $D_P$ | Diffusion of Pak1 in cytoplasm | 5 | a.u. length/s | (0,10) |
| $k_c$ | Activation rate constant of Cdc42 at tips | 1 | | (0,2) |
| $n_{sc}$ | Cooperativity of active Cdc42 self-regulation | 4.5 | | (0,6) |
| $K_{sc}$ | Threshold of Cdc42 activation by Scd1 | 0.545 | a.u | (0,1) |
| $\delta_c$ | Degradation/ removal rate constant of active Cdc42 | 0.3 | /min | (0,1) |
| $k_s$ | Accumulation rate constant of Scd1 at tips | 1.5 | | (0,2) |
| $n_{cs}$ | Cooperativity of Scd1 by active Cdc42 | 0.9 | | (0,4) |
| $K_{cs}$ | Threshold of Scd1 accumulation by active Cdc42 | 1 | a.u. | (0,2) |
| $K_{ps}$ | Threshold of Scd1 inhibition by Pak1 | 0.145 | a.u. | (0,1) |
| $n_{ps}$ | Cooperativity of Scd1 by Pak1 | 3 | | (0,6) |
| $\delta_s$ | Degradation/ removal rate constant of Scd1 | 0.1 | /min | (0,1) |
| $k_p$ | Accumulation rate constant of Pak1 at tips | 2 | | (0,2) |
| $n_{cp}$ | Cooperativity of Pak1 by active Cdc42 | 3 | | (0,6) |
| $K_{cp}$ | Threshold of Pak1 accumulation by active Cdc42 | 0.5 | a.u. | (0,2) |
| $\delta_p$ | Degradation/ removal rate constant of Pak1 | 0.4 | /min | (0,2) |
| a1 | Strength constant of second negative signaling pathway | 0.5 | | |
| $\delta_p'$ | Degradation/ removal rate constant of Pak1 | 0.15 | /min | |
| $k_{pe}$ | Strength constant of endocytosis | 2 | | |
| $n_{pe}$ | Cooperativity of endocytosis/patches by Pak1 | 2 | | |

| | | | |
|---|---|---|---|
| $K_{pe}$ | Threshold of endocytosis by Pak1 | 0.3 | a.u. |

* a.u. is an arbitrary unit of concentration.

** a.u. length is an arbitrary unit of length.

Modified parameters for mutants and Model3 were labeled above their corresponding figures.

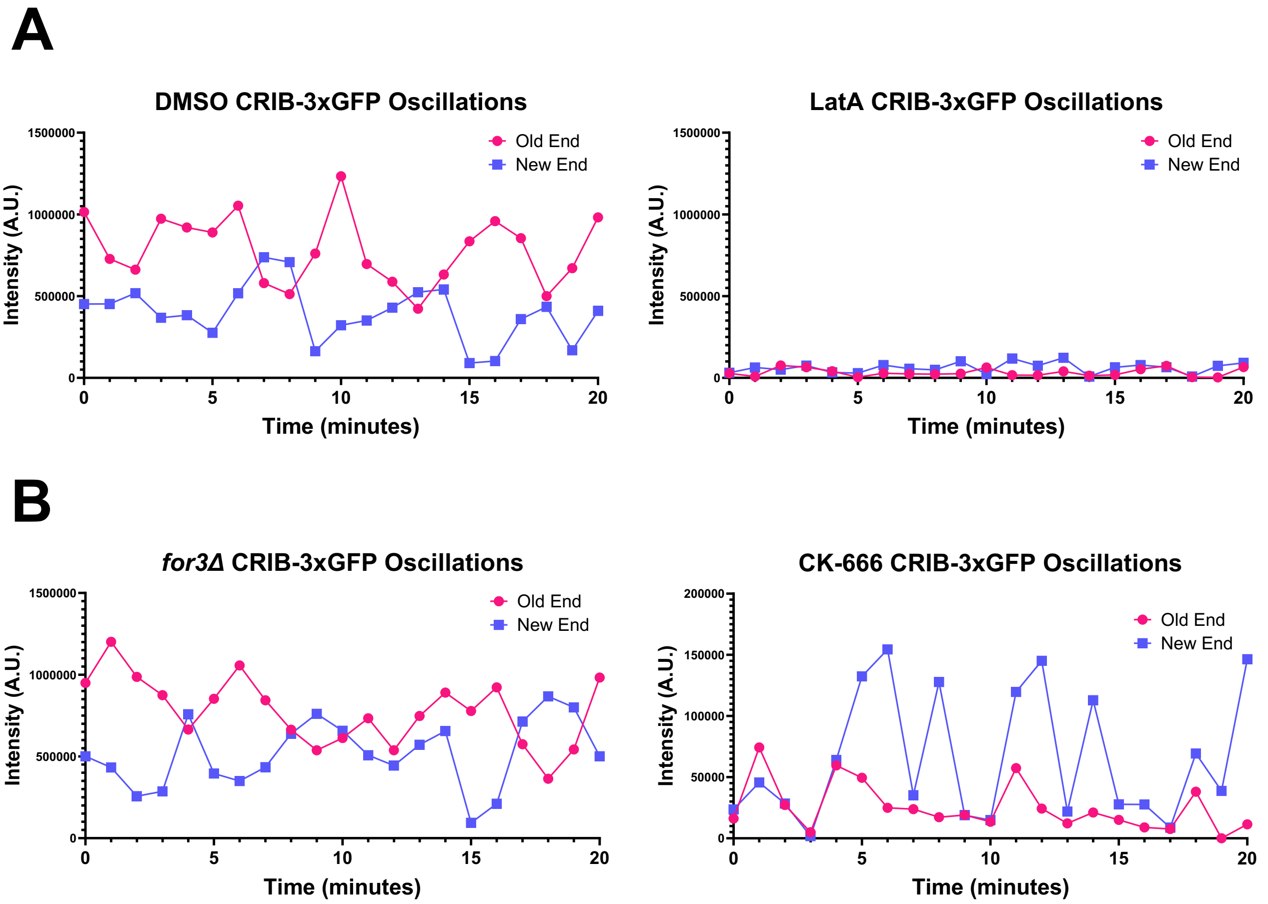

**Supplemental figure 1. Representative CRIB-GFP Oscillations for each condition. A**. Representative CRIB-3xGFP oscillations in DMSO and Lat-A treated cells. **B.** Representative CRIB-3xGFP oscillations in *for3*Δ and CK-666 treated cells.

Supplemental Figure 1

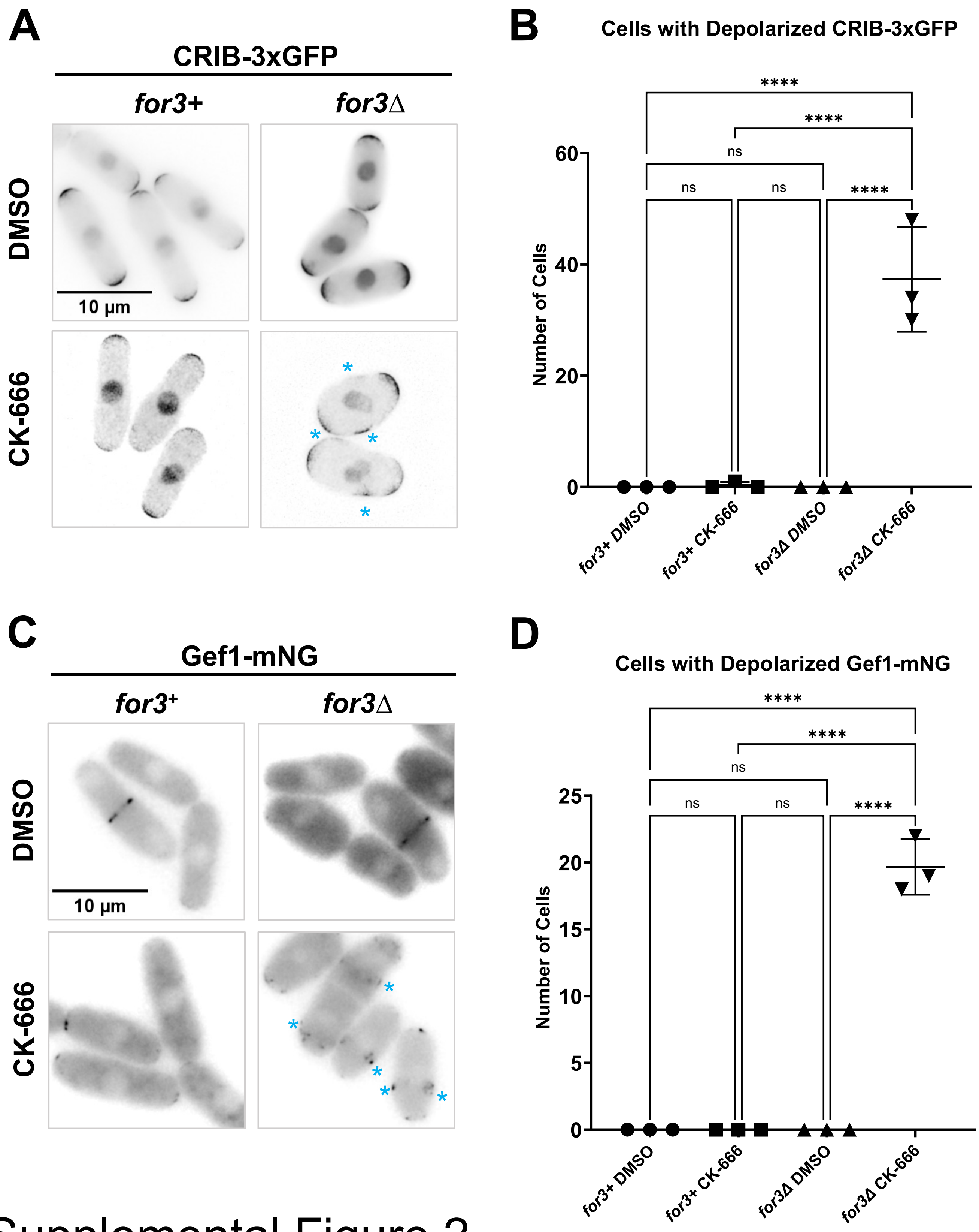

Supplemental Figure 2

**Supplemental figure 2. Loss of both F-actin structures causes a stress response but CK-666 treatment alone does not**. **A.** CRIB-3xGFP localization in *for3+ and for3*Δ cells treated with DMSO and CK-666. **B.** Quantification of the number of cells with depolarized CRIB-3xGFP localization. **C.** Gef1-mNG localization in *for3*+ and *for3*Δ cells treated with DMSO and CK-666. **D.** Quantification of the number of cells with depolarized Gef1-mNG localization. Scale bar, 10μm. n.s., not significant; p-value, ****<0.0001 ,One way ANOVA, followed by Tukey's multiple comparison test.

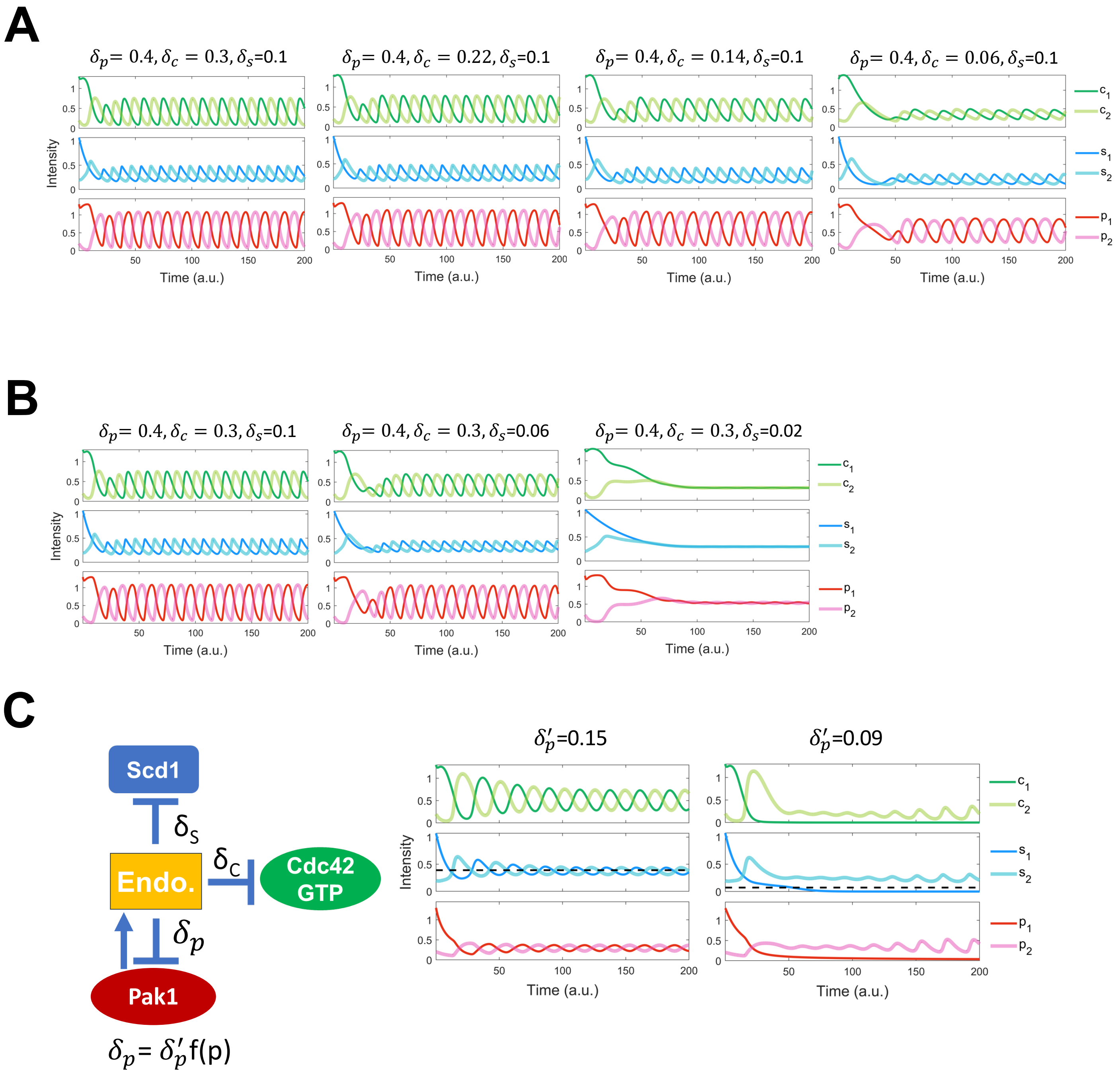

**Supplemental Figure 3. A.** Bipolar dynamics after reducing the removal rate of Cdc42 to 20% of the wild-type removal rate. **B.** Protein dynamics upon reducing the removal rate of Scd1 to 20% of the wild-type rate. **C.**Model 2−1 includes an additional component where Pak1 promotes endocytosis. $\delta_p'$ is a constant, f(p) is the hill function of Pak1. The simulation indicates that upon inhibiting endocytosis, bipolar oscillatory dynamics transition to monopolar dynamics, closely resembling the outcomes of the main Model 2. Details are in the supplementary materials.

### Supplemental Figure 3

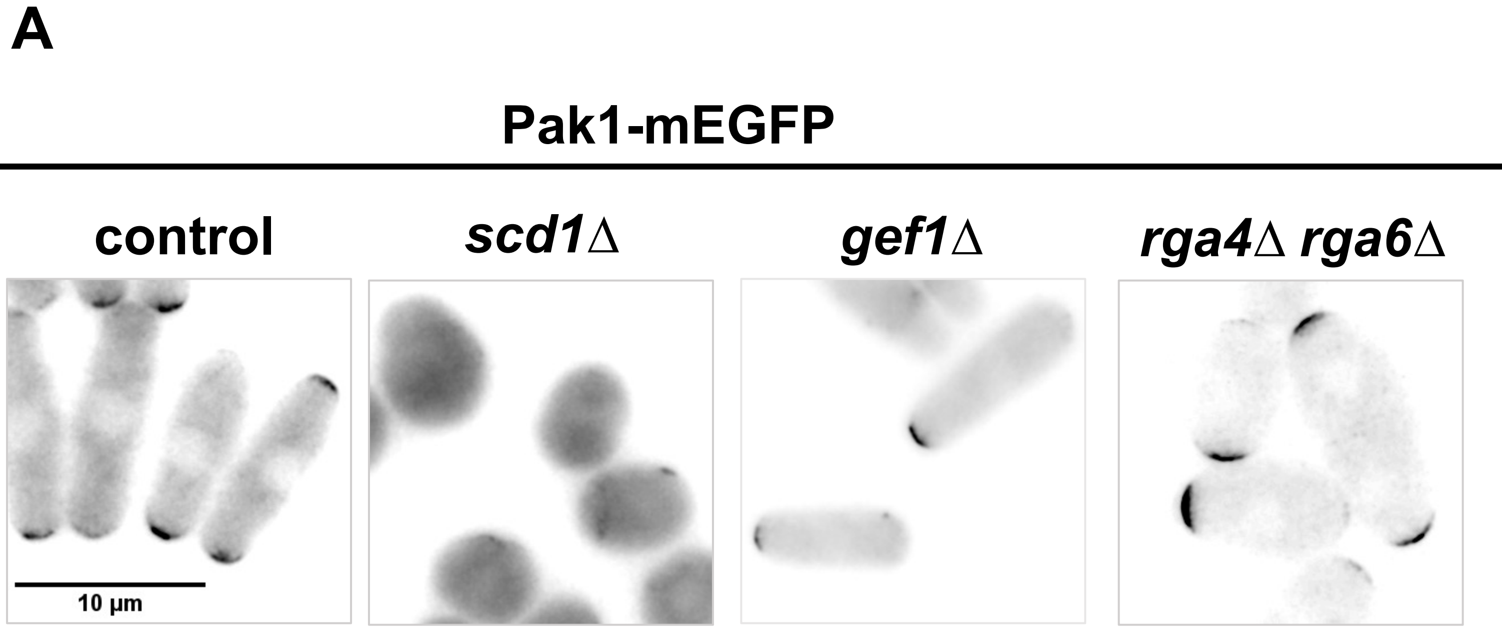

**Supplemental figure 4. Cdc42 regulators Impacts' on Pak1 localization. A.** Pak1-mEGFP localization in control cells, *scd1Δ, gef1Δ ,* and *rga4Δ rga6Δ* mutant cells. Scale bar, 10µm.

Supplemental Figure 4

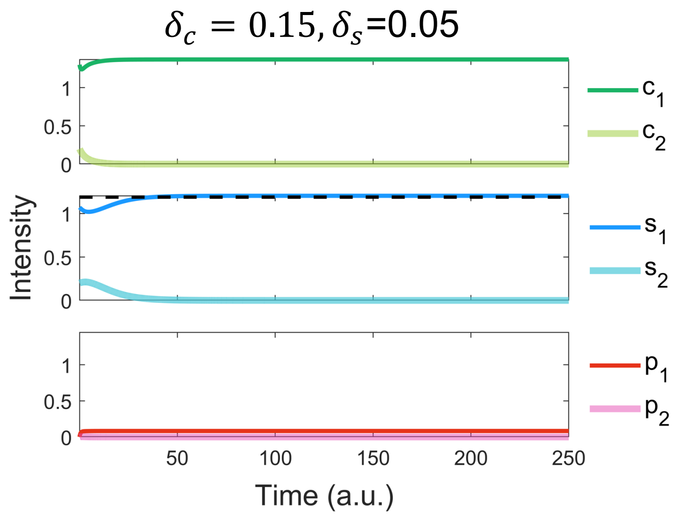

**Supplemental Figure 5. Modeling simulation of localization of Scd1 and Cdc42-GTP in *pak1-ts* cells in Model 2.** Monopolar localization of Scd1 and Cdc42-GTP in the absence of *pak1.*

Supplemental Figure 5

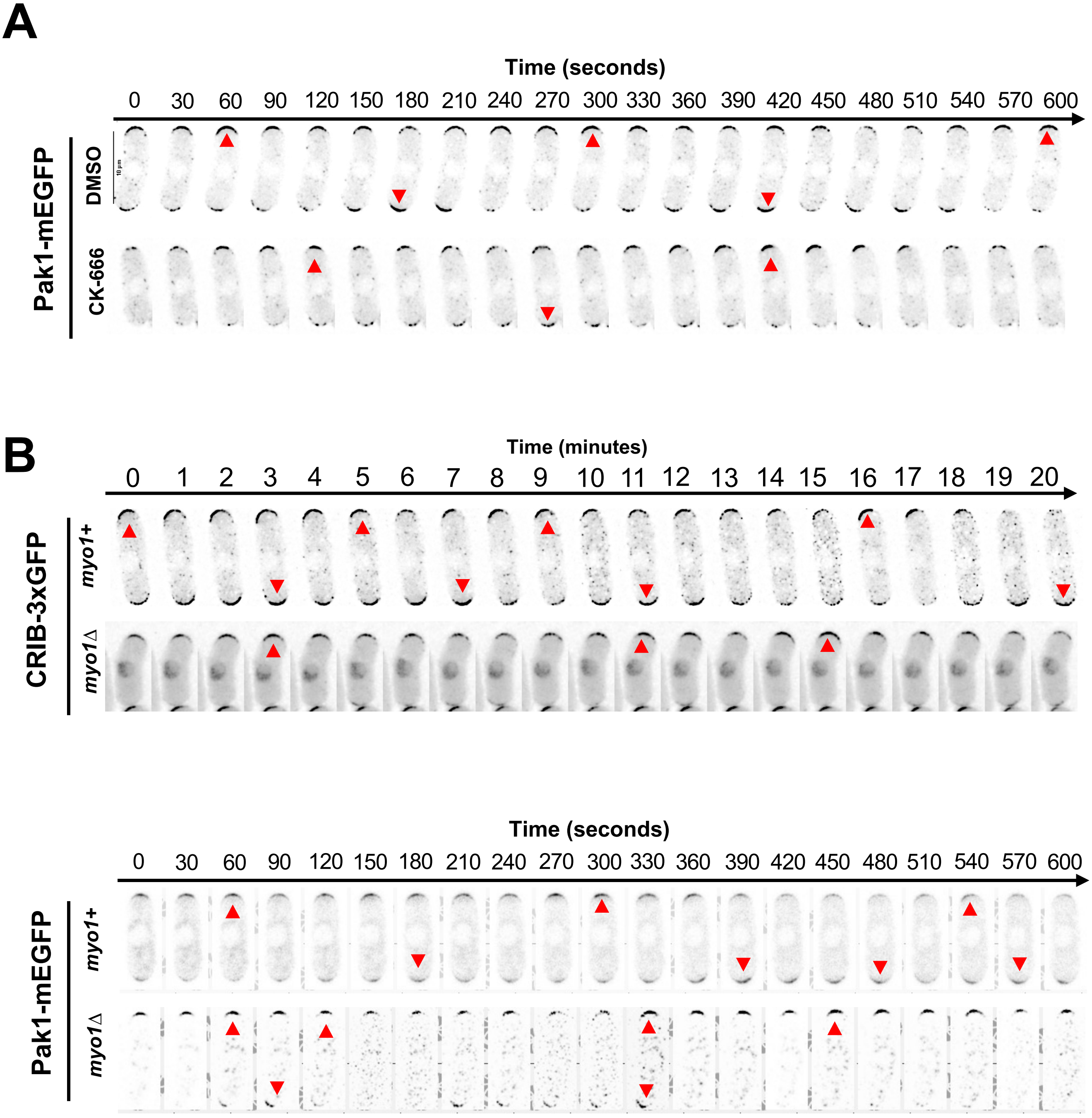

**Supplemental figure 6. Pak1-mEGFP and CRIB-GFP dynamics under different conditions. A.** Pak1-mEGFP dynamics in DMSO and CK-666 treated cells. Red arrows indicate peaks of fluorescence intensity at that cell end. **B.** Dynamics of CRIB-GFP and Pak1-mEGFP were observed in *myo1+* and *myo1*Δ cells. Red arrows indicate peaks of fluorescent intensity at that cell end. Montages clarified and denoised, made in NIS elements. Scale bar 10µm.

### Supplemental Figure 6

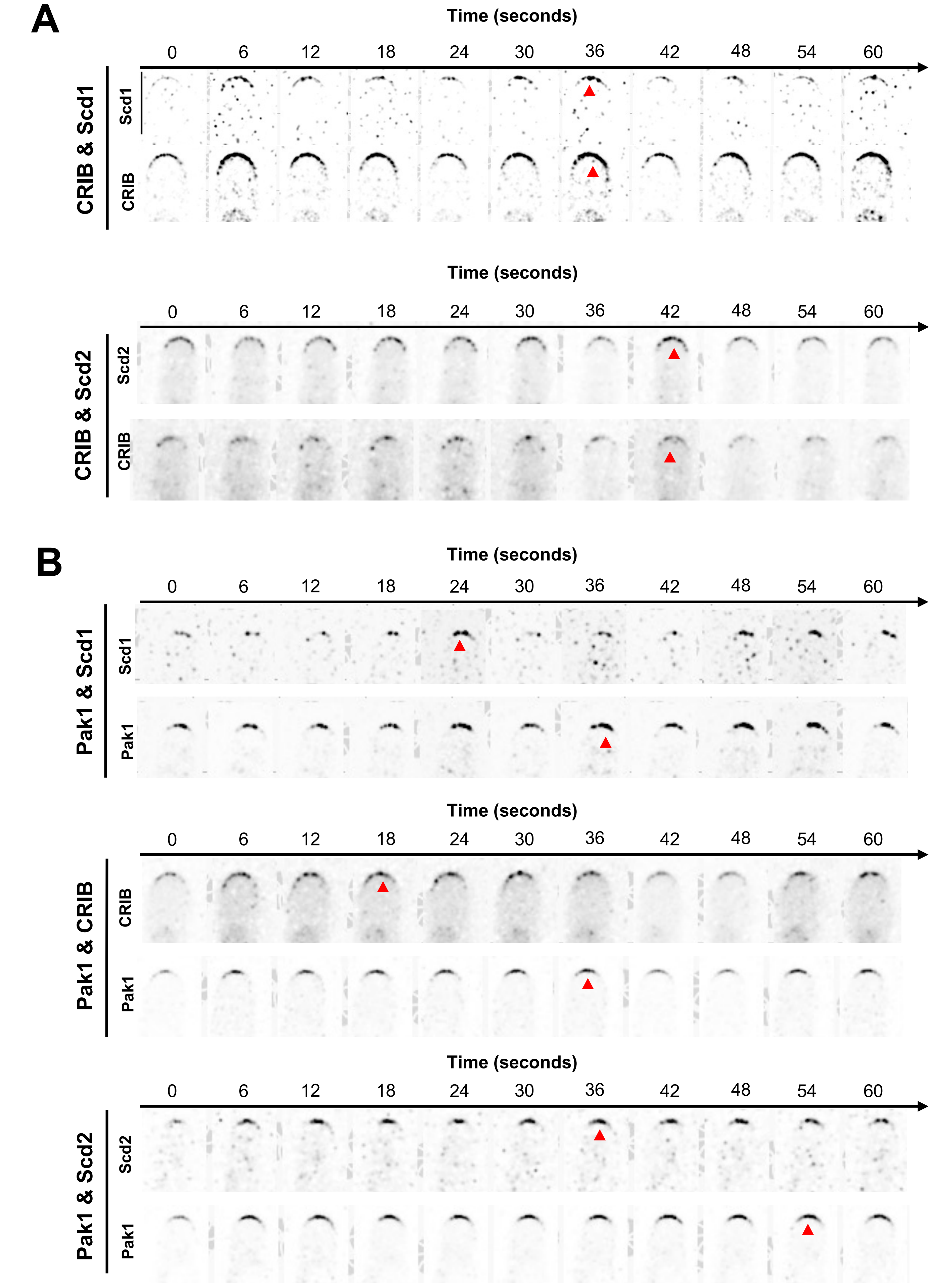

Supplemental Figure 7

**Supplemental figure 7. Montages of phase shift between pairs of polarity proteins. A.** Montage of phase shift between Scd1-tdTomato and Cdc42 activity (CRIB-3xGFP) and between Scd2-mCherry and CRIB-3xGFP). Arrows denote peaks in intensity. **B.** Montage of phase shift between Pak1-mEGFP and the positive feedback proteins: Scd1-tdTomato, active Cdc42 (CRIB-mCherry), and Scd2-mCherry. Arrows indicate peaks in intensity. Montages clarified and denoised, made in NIS elements. Scale bar 5μm.
